# A Modular Layer-by-Layer Nanoparticle Platform for Hematopoietic Progenitor and Stem Cell Targeting

**DOI:** 10.1101/2024.10.30.621186

**Authors:** Tamara G. Dacoba, Namita Nabar, Paula T. Hammond

## Abstract

Effective delivery of drug and gene cargos to hematopoietic stem and progenitor cells (HSPCs) is a major challenge. Current therapeutic strategies in genetic disorders or hematological malignancies are hindered by high costs, low accessibility, and high off-target toxicities. Layer-by-Layer nanoparticles (LbL NPs) are modular systems with tunable surface properties to enable highly specific targeting. In this work, we developed LbL NPs that target HSPCs via antibody functionalization with reduced off-target uptake by circulating myeloid cells. NPs layered with poly(acrylic acid), a bioinert polymer, provided more stealth properties in vivo than other tested bioactive polyanions. The additional conjugation of anti-cKit and anti-CD90 antibodies improved NP uptake by 2-to 3-fold in non-differentiated bone marrow cells in vitro. By contrast, anti-CD105 functionalized NPs showed the highest association to HSPCs in vivo, ranging from 3.0–8.5% in progenitor subpopulations. This LbL NP platform was then adapted to target human HSPC receptors, with similar targeting trends in healthy CD34+ human cells. By contrast, anti-CXCR4 functionalization demonstrated the greatest targeting to human B-cell lymphoma and leukemia cells. Taken together, these results underscore the therapeutic potential of this modular LbL NP platform with the capacity to target HSPCs in a disease-dependent context.

---

Hematopoietic stem and progenitor cells (HSPCs) are the precursors of all blood cell lineages. Any mutations or defects affecting these cells can result in a number of blood diseases, such as hemoglobinopathies (e.g., beta-thalassemia, sickle cell disease), immune disorders, storage and metabolic disorders, or blood cell malignancies (e.g., acute lymphoid leukemia, non-Hodgkin’s lymphoma).^1,2^ HSPCs possess self-renewal and multipotency capacity, making them invaluable therapeutic targets to treat any of these disorders at the source. However, manipulating HSPCs is challenging due to their limited numbers and confinement within the bone marrow niche.^3^

Current clinical interventions for these disorders involve hematopoietic stem cell transplantation, which raises limitations of variable patient responses, limited matched donors, and morbidity and mortality rates due to graft-versus-host disease or cancer relapse.^4^ In monogenic disorders, recent advances in autologous ex vivo HSPC gene editing eliminate the need for matched donors. However, these processes remain highly specialized and costly (over $2M per patient), severely limiting their accessibility to the general population.^5^ Furthermore, these and other therapies in clinical evaluation employ viral vectors or electroporation, which raise safety concerns and still have limited applicability.^1,6,7^ In hematological malignancies, chemotherapeutics or small molecule inhibitors present broad organ biodistribution and poor specificity that lead to serious side effects in healthy tissues.^8,9^ Thus, to ultimately enable broader therapeutic applications in hematological disorders, the development of highly-specific targeted delivery approaches are crucial.^10,11^

To this end, nanoparticles (NPs) can deliver a great variety of drug cargos, while reducing off-target toxicity and improving drug biodistribution.^12,13^ Targeting specificity can be further enhanced by tuning NP surface chemistries through layer-by-layer (LbL) assembly, the electrostatic deposition of alternating polyelectrolytes onto NP substrates.^14,15^ The LbL NP construct’s modularity allows the NP core to be switched based on the cargo of interest, with liposomes,^16–18^ polymeric NPs,^19^ and lipid nanoparticles (LNPs)^20^ amenable to this surface modification. These LbL NPs can be further directed to cells and tissues of interest by adsorbing or conjugating targeting moieties such as peptides^21–23^ or antibodies (Abs)^24^ to their surfaces.

To date, HSPC-targeting strategies have been directed at many different surface markers. For instance, CD45, a universal hematopoietic marker expressed in HSPCs and differentiated immune cells, has been targeted both by antibody-drug conjugates (ADCs) and Ab-conjugated LNPs.^25–27^ The cKit (CD117) receptor, of the tyrosine kinase family, is known for its role in HSPC proliferation, survival, and differentiation.^28,29^ Recently, its targeting via anti-cKit-conjugated LNPs translated into improved gene editing rates in mouse and human HSPCs.^30,31^ However, reported adverse effects for an anti-cKit ADC—including the death of a patient—have raised safety concerns around targeting these receptors, highlighting the need for alternatives.^32^ Another candidate, the CXCR4 (CD184) receptor, is involved in HSPC homing in the bone marrow niche, which has led to the clinical use of its antagonists as HSPC-mobilization agents.^33–35^ Additional HSPC receptors of interest include CD90 (Thy1), a cell surface protein expressed by primitive HSPCs,^36^ and CD105 (endoglin), part of the TGF-β receptor complex and expressed by subsets of HSPCs where it promotes stemness.^37,38^ Both lentiviral vectors and NPs conjugated to Abs against these receptors have shown successful targeting results.^37,39–41^ Despite these advancements, no studies have systematically compared the benefits and limitations of targeting these hematopoietic receptors across both healthy and diseased conditions.

In this work, we evaluate the potential of an LbL NP platform to develop a precision drug delivery system, leveraging the modularity of LbL NPs to compare and assess different surface properties for targeting hematopoietic progenitors. We first explore several surface polymers for their ability to decrease unwanted LbL NP interactions with circulating myeloid cells.^5^ Upon selection of poly(acrylic acid) (PAA) as the stealth layer, we further modify the surface of PAA NPs with HSPC-targeting Abs, systematically comparing various promising candidates: anti-cKit, anti-CD45, anti-CXCR4, anti-CD90, and anti-CD105. In vitro studies in mouse bone marrow cells demonstrate the impact of Ab on LbL NP uptake and trafficking patterns across Abs, with anti-cKit and anti-CD90 improving NP uptake the most. In vivo, anti-CD105 functionalization causes the greatest association to murine HSPCs. To assess the potential translation of these targets, we utilize human CD34+ cells to confirm that anti-human CD105 also increases LbL NP association in vitro. Anti-CXCR4 functionalization confers advantages in disease settings, as it improves LbL NP association in CXCR4-expressing human leukemia and lymphoma lines. Overall, we harness the modularity of LbL NPs to develop a platform with stealth properties that, upon the conjugation of a targeting moiety, can efficiently reach different hematopoietic targets in vitro and in vivo.

## RESULTS

### Outer layer selection impacts uptake by circulating myeloid cells in vivo

Myeloid cells such as granulocytes and monocytes exert a critical function in immune surveillance by clearing foreign bodies, such as NPs, from systemic circulation. To prevent this rapid clearance, we first aimed to identify an LbL NP outer layer that would limit the interactions with these cells. For this, we layered negatively charged fluorescently-labeled liposomes sequentially with the polycation poly-L-arginine (PLR), and a polyanion: hyaluronic acid (HA), poly-L-glutamic acid (PLE), or poly(acrylic acid) (PAA) (Fig. 1A). After each layer deposition, particle size increased by 5–30 nm and charge inverted, yielding final LbL NPs of 100-150 nm diameters, low PDIs, and negative surface charge (Fig. 1B, C). To capture the early interactions that govern NP clearance in vivo, we evaluated LbL NP interactions with circulating immune cells and organ biodistribution 1.5 h after intravenous administration in healthy mice (Fig. 1D and Suppl. Fig. 1). For each of the treated groups, no significant differences were observed in the relative abundances of each cell subpopulation in the blood (Fig. 1E). Among all NP groups, association to circulating leukocytes was comparable (Fig. 1F); however, the distribution of leukocyte subsets within NP-positive populations varied significantly among the tested outer layers (Fig. 1G and Suppl. Fig. 2). For instance, HA NP-positive cells were enriched in granulocytes and monocytes –46.9% and 11.6%, respectively–; while bare NP-positive cells were comprised of 26.5% granulocytes and 2.1% monocytes. In contrast, granulocytes and monocytes represented only 13.9% and 4.5% of PAA NP-positive cells, respectively. Consequently, a higher portion of PAA NP-positive cells were B cells (74.7%), which reflects their abundance in mouse blood (63.5% in control mice) (Fig. 1E, G). Interestingly, 4.3% of HA NP-positive cells were T cells, higher than within any of the other groups (≤ 2.0%), highlighting some degree of specificity of HA NPs toward T cells as well as myeloid cells. On an organ level, all NPs were present in the liver to a similar extent, and PAA NPs resulted in the lowest accumulation in all the other organs profiled (spleen, lungs, heart, and kidneys) (Fig. 1H and Suppl. Fig. 3). Collectively, PAA’s ability to reduce NP association with myeloid cells and lower organ accumulation led to the selection of this polymer as the stealth outer layer of the LbL NP platform used in this work, creating the basis for Ab conjugation to drive LbL NPs to the desired HSPC while minimizing engagement with nontargeted myeloid cells.

**Fig. 1.**
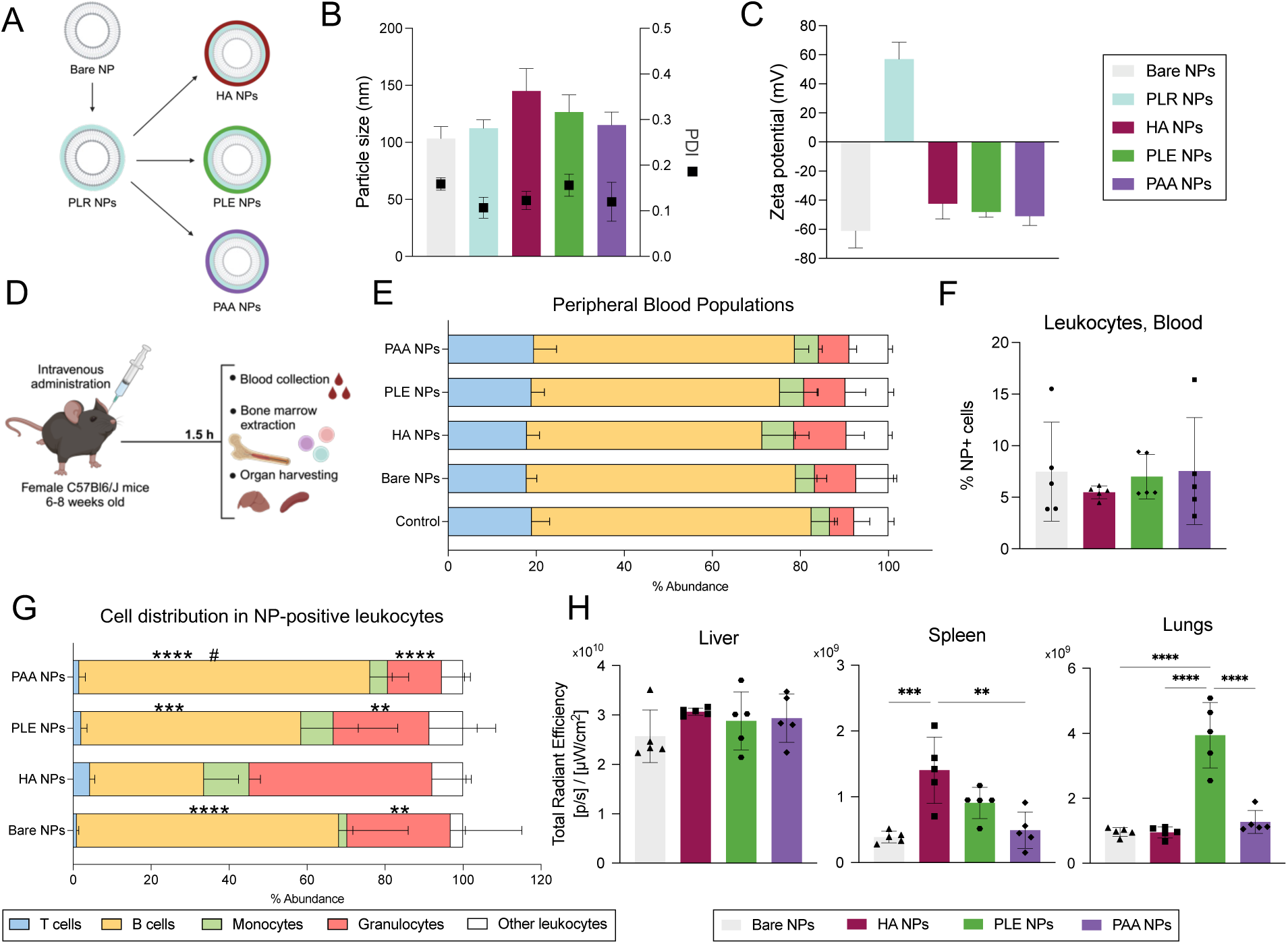
Biodistribution of LbL NPs with HA, PLE, and PAA as outer layers in circulating blood cells and major organs. (**A**) Schematic of the layering assembly, starting with a liposome (bare NP), then layering with the polycation PLR, and finally with three different polyanions: HA, PLE, or PAA. (**B, C**) Average particle size (Z-average), polydispersity index (PDI) and ζ-potential of the different LbL NPs. (**D**) Timeline of in vivo study, in which NPs are intravenously administered to mice, and after 1.5 h the blood and organs are analyzed. (**E**) Blood cell composition in the different treatment groups, in terms of B cells, T cells, monocytes, granulocytes, and other leukocytes. (**F**) LbL NP association with all CD45+ leukocytes for the different treatment groups; bars represent the mean, and each dot one biological replicate. (**G**) Blood cell distribution of these NP-positive cells for the different cell subtypes. Statistical analysis is a two-way ANOVA with Tukey correction (* are differences compared to HA NP group and # to PLE NP group; *p<0.05, **p<0.01, ***p<0.005, ****p<0.001); n=5 mice per group. (**H**) Radiant efficiency measurements obtained via IVIS of the fluorophore (Cy5)-tagged LbL NPs in liver, spleen, and lungs; bars represent the mean, and each dot represents one biological replicate; n=5 mice per group. Statistical analysis is a one-way ANOVA with Tukey correction (**p<0.01, ****p<0.001).

### Layer-by-layer nanoparticle functionalization enables modular antibody conjugation

To covalently attach Abs to PAA NPs, we modified approximately 2.5% of the carboxylic acid groups of PAA with a methyltetrazine amine (mTz) handle, to facilitate an orthogonal reaction with trans-cyclooctene (TCO)-modified Abs (Suppl. Fig. 4). We selected five different HSPC targets: cKit, CD45, CXCR4, CD90, and CD105, and generated a library of anti-mouse Ab-functionalized PAA NPs (Ab–PAA NPs) with a non-targeting isotype control (ISOT–PAA NP) (Fig. 2A). In all cases, Abs were partially reduced to expose a low number of thiols and subsequently functionalized via maleimide-thiol chemistry with a bifunctional PEG spacer (maleimide–PEG3– TCO) (Fig. 2B). All Abs had an average of 1-2 linkers (Fig. 2C). To assemble the targeted delivery system, we layered PLR NPs with a 50/50 blend of PAA and mTz-PAA, resulting in ∼120 nm particle sizes, with low PDI, and negative surface charge (Fig. 2D, E). A high Ab conjugation yield (>80%) was achieved across all groups (Fig. 2F). Cryo transmission electron microscopy images of PAA NPs and cKit–PAA NPs confirmed the NP structure was maintained after functionalization (Fig. 2G and Suppl. Fig 5A). The association of Abs to the surface of PAA NPs was further validated through staining with 6-nm gold NPs functionalized with an Ab binding fragment.^42,43^ Using polystyrene NPs as the core, gold NPs were detected on the surface of Ab–PAA NPs, in comparison to PAA NPs (Fig. 2H and Suppl. Fig 5B). By analyzing the number of gold NPs per PAA NP, we calculated that approximately 45 Abs are conjugated per NP. Overall, through optimized surface chemistry and conjugation, we have developed a highly modular Ab–LbL NP template to evaluate a broad library of HSPC-targeting Abs.

**Fig. 2.**
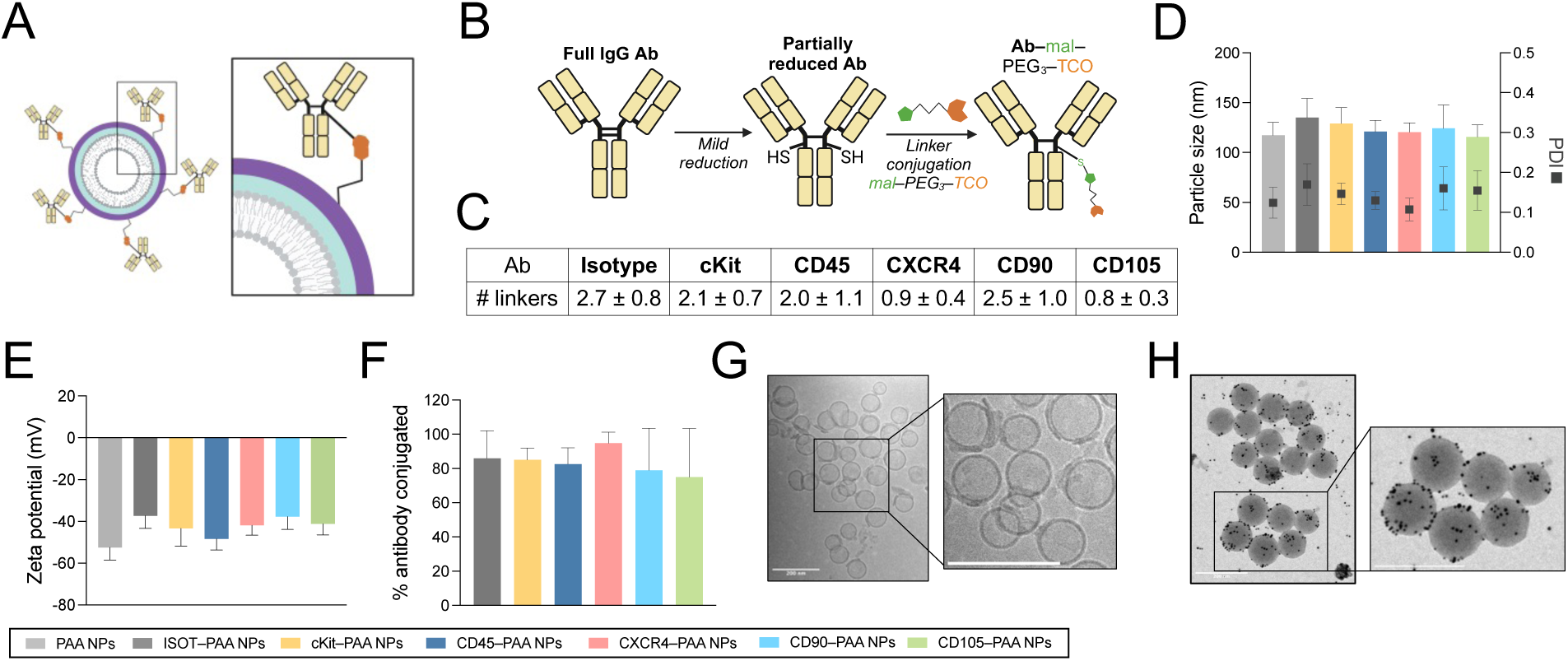
Preparation of a modular Ab–PAA NP platform. (**A**) Schematic of an Ab–LbL NP, zooming into the Ab conjugated through the hinge region to the chemical handles presented on the outer polymer layer. (**B**) Ab modification reaction, where a full IgG Ab is reduced to free the thiol groups in the hinge region and then reacted to a bifunctional linker. (**C**) Table summarizing the number of linkers conjugated to each Ab (n≥3 replicates). (**D, E**) Average particle size (Z-average), polydispersity index (PDI), and ζ-potential of the library of Ab–PAA NPs. (**F**) Yield of the conjugation reaction after purification of the five Abs and control isotype. Bars represent mean ± standard deviation (n≥3 replicates). (**G**) Cryo transmission electron microscopy images of cKit–PAA NPs. (**H**) Transmission electron microscopy images of cKit–PAA NPs (with a 100-nm polystyrene core) stained with 6-nm gold NPs functionalized with an Ab binding fragment. Scale bars represent 200 nm.

### Targeting antibodies dictate in vitro uptake and trafficking in hematopoietic progenitors

To better understand the interactions of the selected Abs with target cells, we characterized the expression of their corresponding receptors in freshly harvested mouse whole bone marrow (WBM) cells and in lineage-negative (Lin-) cells (Fig. 3A). Lin-cells are WBM cells that have been depleted of lineage-committed cells such as T cells, B cells, monocytes/macrophages, granulocytes, and erythrocytes. In both populations, more than 90% of cells expressed CD45, as expected for this universal hematopoietic marker.^26^ The other receptors were more highly expressed in Lin-cells than WBM, confirming their specificity for progenitor-like populations (Fig. 3B).

**Fig. 3.**
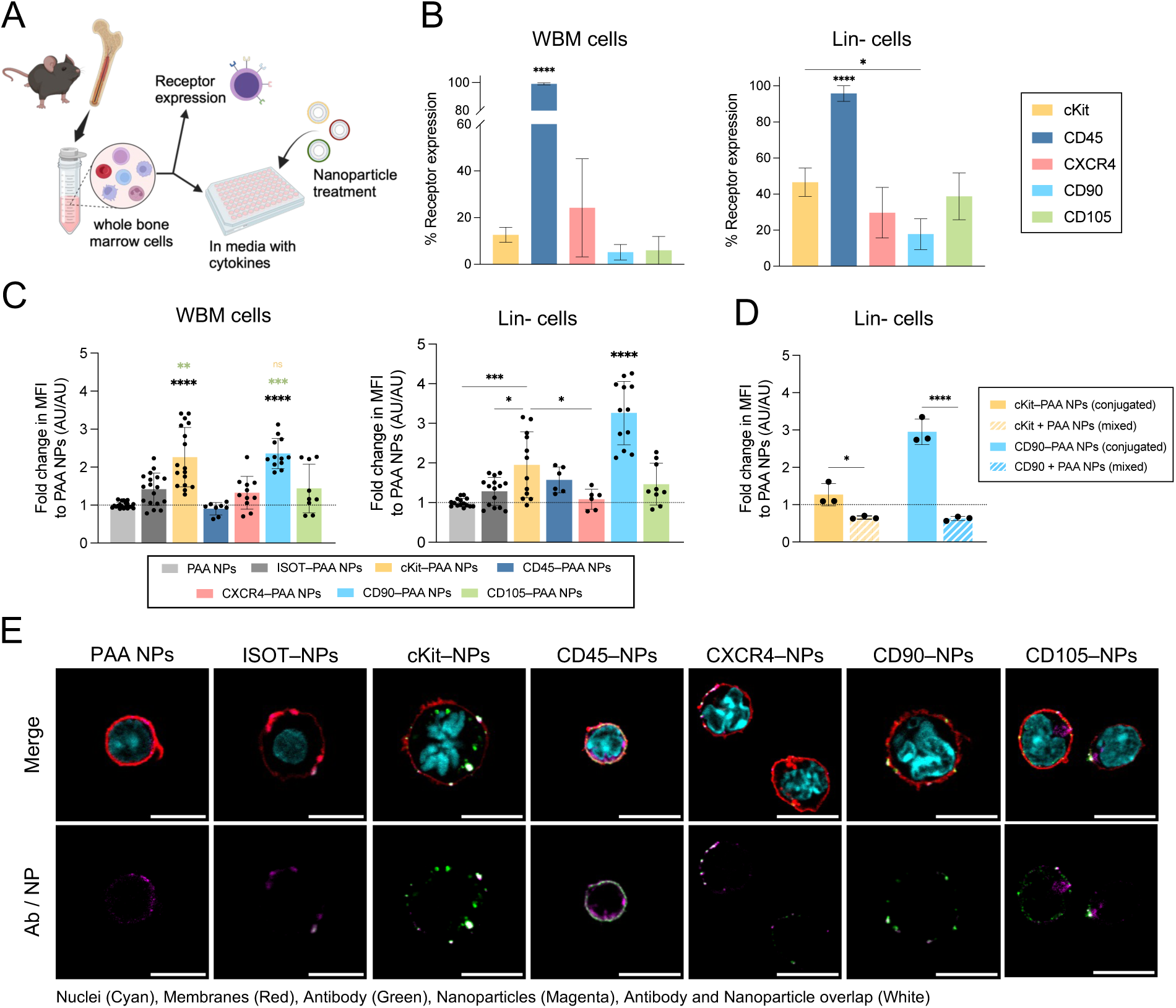
Evaluation of Ab–PAA NPs in bone marrow cells. (**A**) Bone marrow cells were freshly harvested to measure receptor expression, or to evaluate Ab–LbL NP association and trafficking after expansion in cell culture media with cytokines. (**B**) Expression of receptors for each targeting Ab in WBM cells and Lin-cells. (**C**) Median fluorescence intensity (MFI) values for all groups after 1.5 h of incubation with WBM cells or Lin-cells, normalized to PAA NP. Statistical analysis is a one-way ANOVA with Tukey correction (*p<0.05, ***p<0.005, ****p<0.001). Each point is one technical replicate (≥3 biological replicates, each with 3 technical replicates). (**D**) Median fluorescence intensity (MFI) values after 1.5 h of incubation of Ab–PAA NPs or PAA NPs with unconjugated Ab in Lin-cells, normalized to PAA NP. (**E**) Representative confocal images of WBM cells that were treated with Ab–PAA NPs for 1.5 h. Top images represent the merge of all the color stains, and the bottom images show the NP core (Cy5-tagged lipid) and Ab (secondary antibody) stains. Scale bars represent 10 µm.

Next, we evaluated the association of PAA NPs and Ab–PAA NPs to WBM and Lin-cells after short incubation times (0.5 and 1.5 h) to capture Ab-mediated uptake. Flow cytometry showed that anti-cKit and anti-CD90 functionalization yielded the highest NP association (Fig. 3C and Suppl. Fig 6A, B). Indeed, NP association to Lin-cells at 1.5 h increased by 2.0- and 3.3-fold with cKit– and CD90–PAA NPs, in comparison to PAA NPs (Fig. 3C). We further confirmed this effect was driven by the conjugation of Abs to PAA NPs, and not solely by the presence of non-functionalized Abs mixed with PAA NPs (Fig. 3D and Suppl. Fig. 6C).

We then characterized the trafficking of Ab–PAA NPs in WBM cells by confocal microscopy, tracking both the NP core (via Cy5-tagged lipid) and the conjugated Ab (via secondary Ab staining) at 1.5 h of incubation. Both untargeted PAA NPs and ISOT–PAA NPs exhibited low NP-cell interactions (Fig. 3E). For cKit– and CD105–PAA NPs, both NPs and Abs were localized inside the cells and associated to cell membranes. For CD45–PAA NPs, a strong Ab and NP signal was detected on the surface of WBM cells, with no NP internalization, correlating with the high CD45 receptor expression on hematopoietic cells (Fig. 3B). Both CXCR4– and CD90–PAA NPs presented similar co-localization patterns with cell membranes, although flow cytometry indicated a stronger association to WBM cells for CD90–PAA NPs (Fig. 3E). Overall, functionalization of PAA NPs with HSPC-targeting Abs induces significant changes in the uptake and trafficking patterns within bone marrow populations, with anti-cKit and anti-CD90 promoting a more effective NP targeting in vitro compared to anti-CD45, anti-CXCR4, or anti-CD105.

### Anti-CD105 functionalization improves HSPC-targeting capacity of PAA NPs in vivo

Next, we investigated the in vivo targeting capacity of the Ab–PAA NP platform. In this study, Ab– PAA NPs were injected intravenously, and NP uptake was analyzed in bone marrow, blood, and major organs after 1.5 h (Fig. 4A). In bone marrow, we identified key hematopoietic subpopulations via flow cytometry (Fig. 4B and Suppl. Fig. 7). Within Lin-cells, we evaluated Lin-cKit+ Sca-1+ (LSK) cells, multipotent progenitor (MPP) subsets, and long-term hematopoietic stem cells (HSC) subpopulations. Although long-term HSCs are the only cell population with self-renewing and long-term multilineage reconstitution properties, MPP subsets retain some short-to long-term reconstitution capacity, and will produce any hematopoietic lineages –in the case of MPPs– or their committed cell lineages –for granulocyte/monocyte (G/M) MPPs and megakaryocyte/erythrocyte (Mk/E) MPPs.^44^ Moreover, the HSPC compartment is increasingly recognized as dynamic and less strictly hierarchical than traditionally modeled, with both long- and short-term progenitors playing crucial roles in contexts such as hematopoietic reconstitution.^45^ Therefore, targeting any of this diverse set of hematopoietic progenitors is expected to yield therapeutic benefits in hematopoietic disorders.

**Fig. 4.**
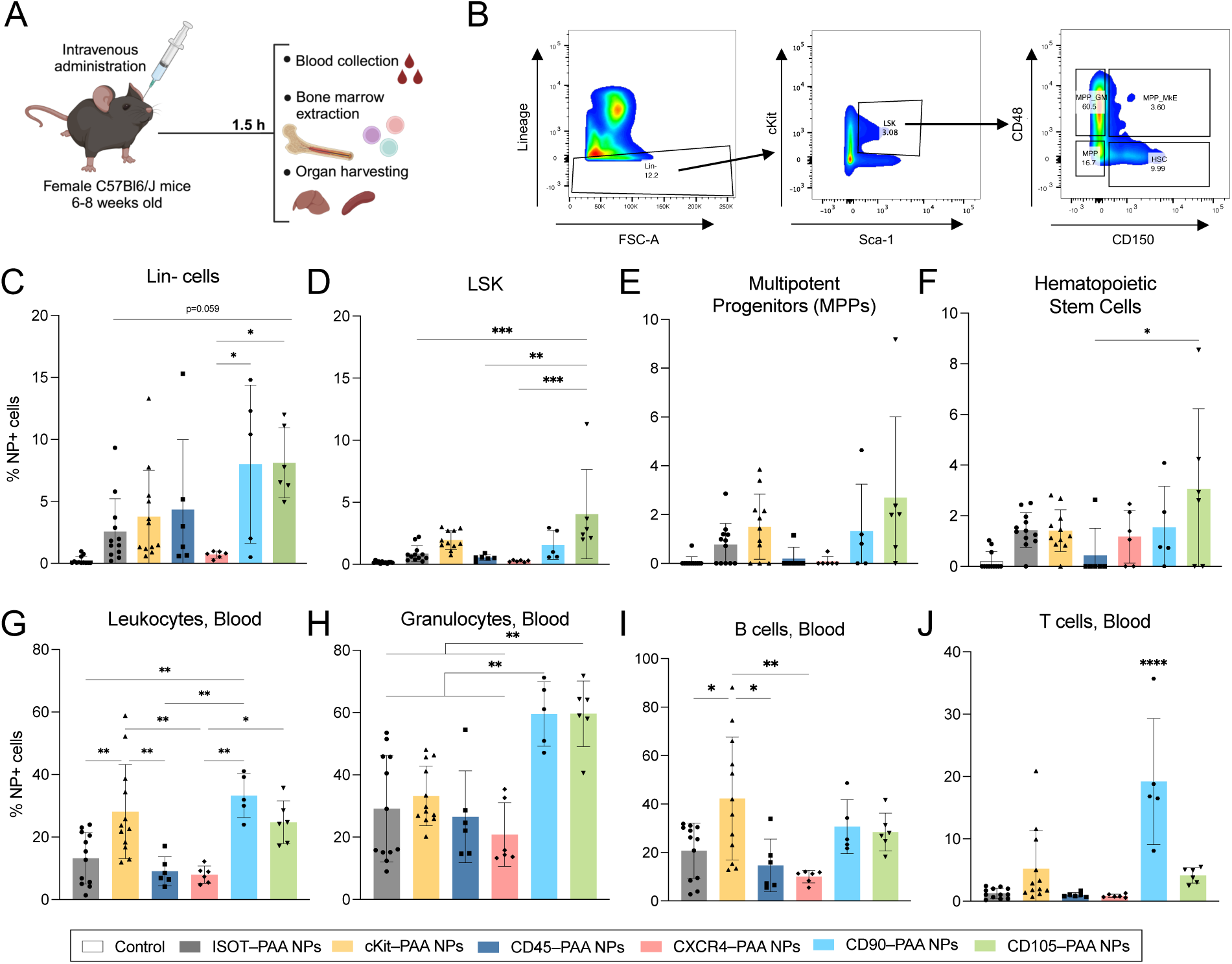
Ab–PAA NPs improve NP targeting capacity of hematopoietic progenitors. (**A**) Timeline of the in vivo study, where Ab–PAA NPs are intravenously administered to mice, and blood, bone marrow, and organs are analyzed after 1.5 h. (**B**) Flow gating strategy for Lin-progenitors, Lin-cKit+ Sca-1+ cells (LSK), different multipotent progenitors (MPPs), and hematopoietic stem cells (HSCs). Ab–PAA NP association to bone marrow cell populations as (**C**) Lin-cells, (**D**) LSK cells, (**E**) MPPs, and (**F**) HSCs. Ab–PAA NP association to blood-circulating (**G**) leukocytes, (**H**) granulocytes, (**I**) B cells, and (**J**) T cells. Statistical analysis is a one-way ANOVA with Tukey correction among Ab–PAA NP groups (*p<0.05, **p<0.01, ***p<0.005, ****p<0.001). Bars represent mean, with standard deviation, and each dot is one animal replicate (n ≥ 5 per group).

CD45– and CXCR4–PAA NPs showed minimal improvement over ISOT–PAA NPs (Fig. 4C-F and Suppl. Fig. 8A-C). By contrast, functionalization with anti-cKit, anti-CD90, and anti-CD105 boosted in vivo targeting of bone marrow cells; in particular, CD90– and CD105–PAA NPs showed the highest targeting to WBM and Lin-cells (Fig. 4C-F and Suppl. Fig. 8A-C). Both CD90– and cKit–PAA NPs showed higher association to MPPs and HSCs than ISOT–PAA NPs, albeit not to a statistically significant degree (Fig. 4E, F and Suppl. Fig. 8B, C). CD105–PAA NPs demonstrated significantly improved uptake by all MPPs and HSCs compared to the other Ab–PAA NPs, associating with 2.7% of MPPs, 3.8% of G/M MPPs, 8.5% of Mk/E MPPs, and 3.0% of HSCs (Fig. 4E, F and Suppl. Fig. 8B, C). These modest uptake levels are highly promising, as previous studies have shown that achieving gene editing in even a small proportion of HSPCs (<10%) can effectively ameliorate disease phenotypes in conditions such as thalassemia.^46,47^

In this study, the cKit receptor expression was significantly reduced in mice that received cKit– PAA NPs, due to the rapid internalization of the receptor and the short analysis timepoint.^48^ The gating strategy for LSK cells in this group was adjusted accordingly (Suppl. Fig. 7C, D). Besides, the low frequencies of HSPCs (0.01–0.03% of WBM cells)^49^ limits the numbers profiled, explaining the variability seen between some animals. Overall, when comparing in vitro and in vivo studies, we replicated targeting improvements with anti-cKit and anti-CD90 Abs and identified an outstanding in vivo targeting capacity with anti-CD105, an effect that was not seen in vitro.

In parallel, we analyzed the interactions of different Ab–PAA NPs with circulating immune cells via flow cytometry, considering that the functionalization of PAA NPs with Abs might also influence these interaction patterns. Similarly to bone marrow cells, cKit–, CD90–, and CD105–PAA NPs demonstrated increased affinity for circulating leukocytes (Fig. 4G). Specifically, cKit–PAA NPs were highly taken up by circulating B cells, while CD90– and CD105–PAA NPs were strongly associated with granulocytes and monocytes (Fig. 4H, I and Suppl. Fig. 9D). CD90–PAA NPs also significantly improved T cell uptake, with 19.2% NP-positive T cells, in contrast to less than 5% for the other groups (Fig. 4J). Ex vivo organ imaging indicated higher spleen and liver NP accumulation in the cKit–, CD90– and CD105–PAA NP treated groups (Suppl. Fig 9).

### Single-targeting is comparable to dual-targeting strategy in vivo

Given anti-CD105’s notable targeting in vivo, we sought to evaluate whether HSPC uptake could be further improved with a dual-targeting strategy, conjugating pairs of high-performing Ab candidates to PAA NPs. We prepared CD105/cKit–PAA NPs, CD105/CD90–PAA NPs, and cKit/CD90–PAA NPs, and compared NP uptake to that of CD105–PAA NPs (Suppl. Fig. 10A–D). In vitro, the dual Ab combinations at different ratios of Abs did not improve uptake over PAA NPs in WBM or Lin-cells (Suppl. Fig. 10E, F). Nevertheless, considering that previous in vitro evaluation had overlooked the targeting of CD105–PAA NPs, we proceeded to evaluate the dual-targeted strategies in vivo. Mouse bone marrow, blood, and major organs were analyzed 1.5 h after intravenous administration of the dual-targeted Ab–PAA NPs at a 1:1 Ab ratio (Fig. 5A). Flow cytometry revealed no significant differences in uptake among groups in WBM, Lin-, or LSK cells, and the NP association of the single- and dual-targeted strategies was comparable in both progenitor subsets and blood-circulating immune cells (Fig. 5B-E and Suppl. Fig. 11). Notably, the B-cell and T-cell tropism of anti-cKit and anti-CD90 functionalization was retained within these combinations (Fig. 5F, G). Differences in NP accumulation in the spleen were also observed (Suppl. Fig. 12). Ultimately, the single-targeted strategy with anti-CD105 was as effective as the combination of Abs against several receptors. Each of the selected Abs might be driving HSPC targeting via distinct mechanisms or by aiding in transport and accumulation in different tissues. Thus, combining them may not further enhance the targeting since their effects might not overlap or complement each other in a way that boosts overall efficacy.

**Fig. 5.**
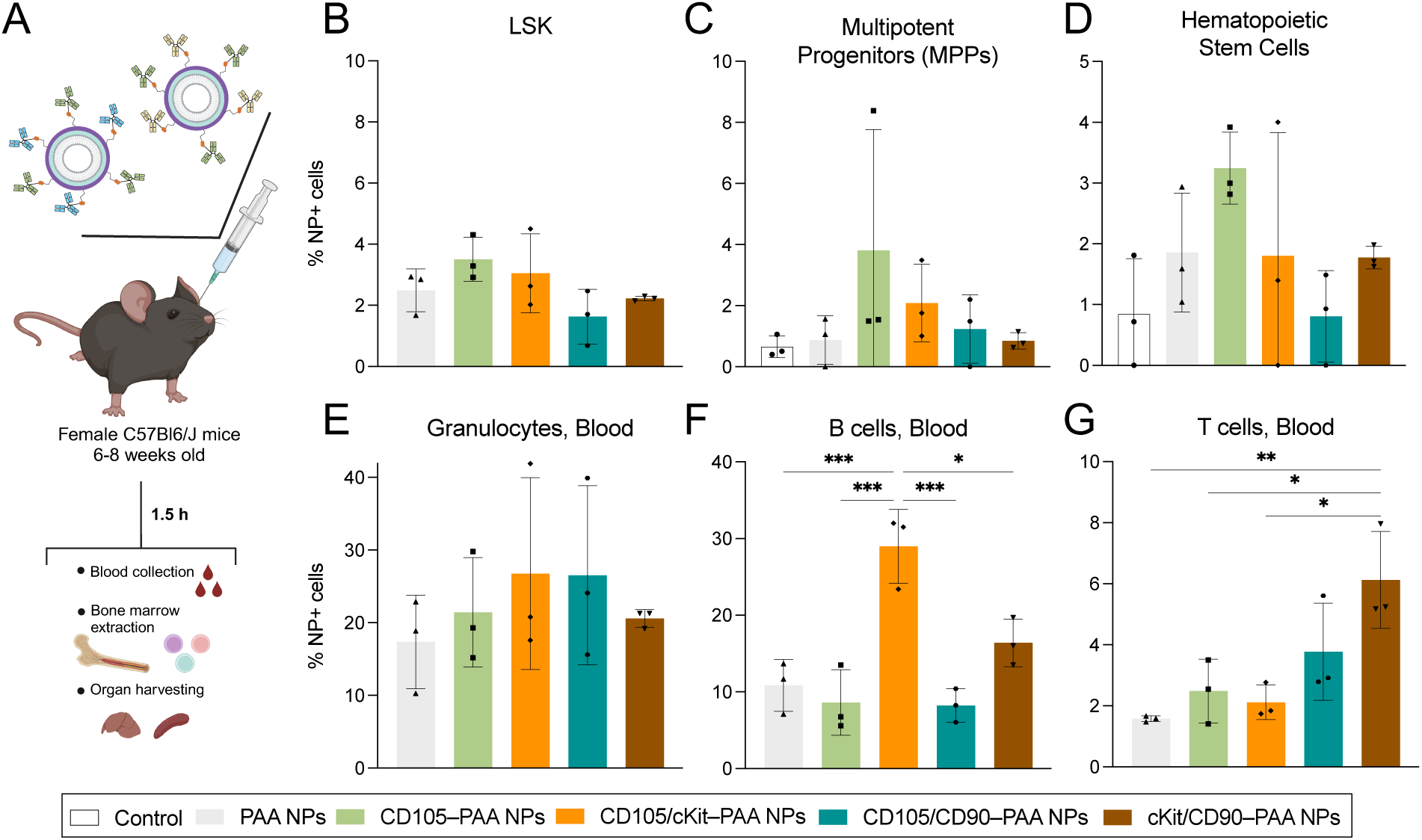
Evaluation of the dual targeted Ab–PAA NP strategy. (**A**) Timeline of in vivo study with dual-targeted Ab–PAA NPs. NP association of the dual targeted Ab–PAA NPs to (**B**) LSK, (**C**) MPPs, and (**D**) HSCs. Dual targeted Ab–PAA NP association to blood-circulating (**E**) granulocytes, (**F**) B cells, and (**G**) T cells. Statistical analysis is a one-way ANOVA with Tukey correction among Ab–PAA NP groups (*p<0.05, **p<0.01, ***p<0.005). Bars represent mean, with standard deviation, and each dot is one animal replicate (n = 3 per group).

### Targeting capacity is translated to human hematopoietic progenitors

Encouraged by the targeting capacity observed in murine models, we explored the translation of this Ab–PAA NP platform to target human HSPCs. For this, Abs against human cKit, CD45, CXCR4, CD90, and CD105 receptors were modified and conjugated to PAA NPs as previously described, with no notable changes to the physicochemical properties of the resulting Ab–PAA NPs (Suppl. Fig. 13). We used healthy human CD34+ cells, as this surface marker is well-established for enriching HSPCs in transplantation and gene editing applications.^45^ To account for patient heterogeneity, cells were sourced from individuals with diverse genetic backgrounds. Human HSPCs were treated with Ab–PAA NPs for 1.5 h, and then NP uptake was analyzed via flow cytometry. While some variability among samples likely reflects differences between donors, CD105–PAA NPs still achieved the highest association to human HSPCs in vitro (Fig. 6A).

**Fig. 6.**
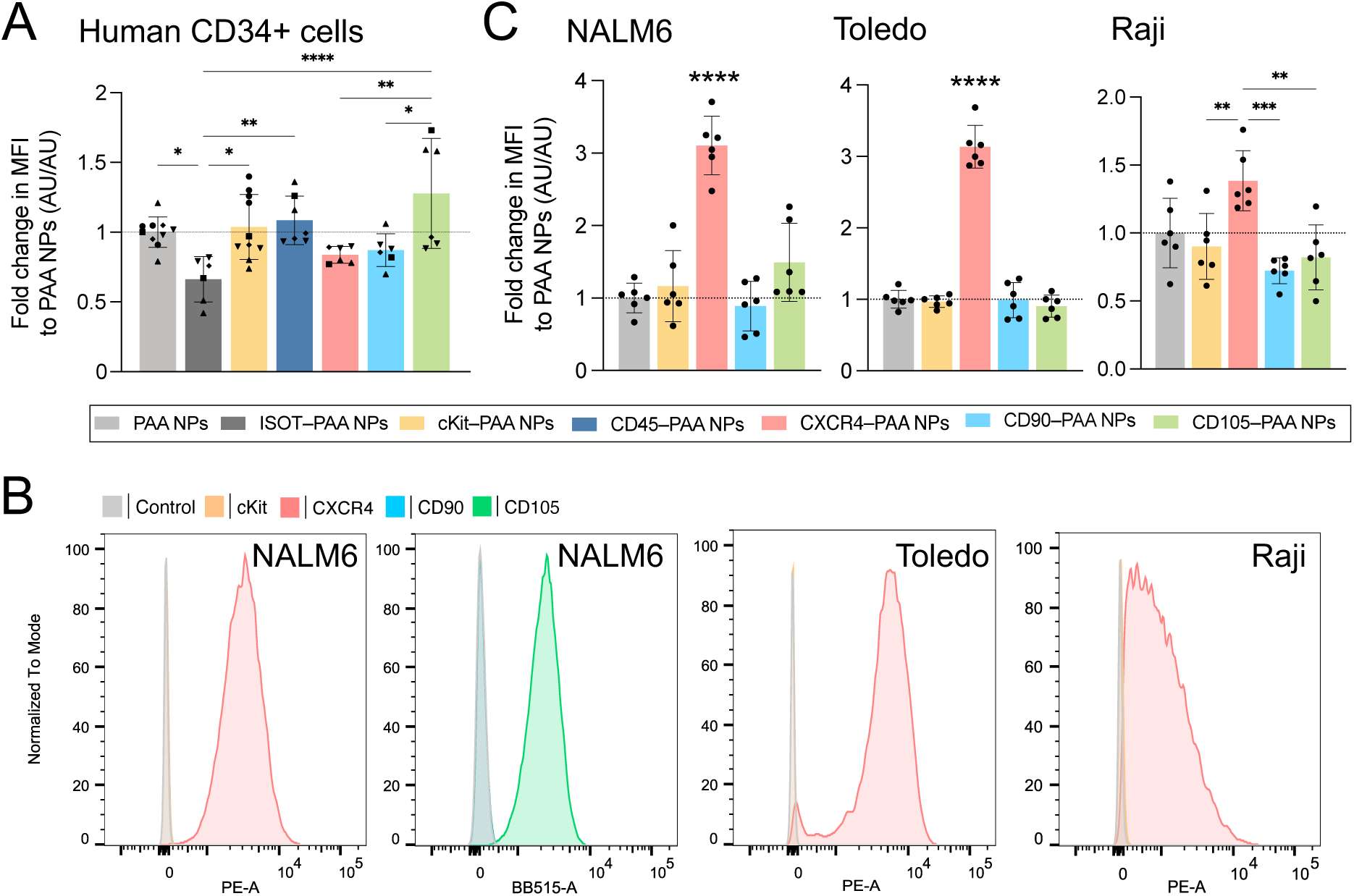
Evaluation of Ab–PAA NP targeting human hematopoietic progenitors. (**A**) Median fluorescence intensity (MFI) values for all Ab–PAA NP groups after 1.5 h of incubation in human CD34+ cells, normalized to PAA NPs. (**B**) Expression of surface receptors for cKit, CXCR4, CD90, CD105 on NALM6 cells, Toledo cells, and Raji cells. (**C**) Median fluorescence intensity (MFI) values after 1.5 h of incubation for anti-human Ab–PAA NPs, normalized to PAA NPs, in NALM6 cells, Raji cells, and Toledo cells. Statistical analysis is a one-way ANOVA with Tukey correction (**p<0.01, ***p<0.005, ****p<0.001). Bars represent mean with standard deviation. For (A) each shape represents a different donor (4 donors, 1-3 technical replicates). For (C) each dot is one technical replicate (2 biological replicates with 3 technical replicates each).

Last, we evaluated the capacity of the Ab–PAA NP platform to target different B-cell derived malignancies: B-cell acute lymphoblastic leukemia (ALL), diffuse large B-cell lymphoma, and Burkitt lymphoma, evaluated in NALM6, Toledo, and Raji cell lines, respectively. Surface staining revealed the most abundant receptor in these three cell lines was CXCR4, while only the CD105 receptor was highly expressed in NALM6 cells (Fig. 6B and Suppl. Fig. 14A). In line with this, CXCR4–PAA NPs presented a 3.1- and 2.1-fold increase in association to NALM6 cells after 1.5 and 4 h of incubation, respectively, in comparison to PAA NPs. CD105–PAA NPs presented a more modest increase, by 1.5- and 1.3-fold (Fig. 6C and Suppl. Fig. 14B). Similarly, anti-CXCR4 functionalization boosted NP uptake in Toledo cells by 3.1- and 2.5-fold, and in Raji cells by 1.4- and 1.5-fold (Fig. 6C and Suppl. Fig. 14B).

## DISCUSSION

In this work, we present a modular NP delivery platform that enables precise cell targeting through rationally designed surface chemistries. By leveraging LbL approaches to decorate NPs with adsorbed polyanions and conjugated Abs, this platform provides the capacity to systematically compare the impact of each component on NP performance and clinical utility.

To design precision targeted NPs in vivo, we first aimed to reduce NP interactions with circulating granulocytes and monocytes, which are known to rapidly clear systemically administered nanocarriers.^49,50^ The outer polymer on LbL NPs can strongly dictate interaction with surrounding cells, as shown in prior studies evaluating LbL NP uptake in cancerous and non-cancerous cells.^51–53^ PAA has shown to decrease NP interactions in vitro with macrophages and monocytes, while HA and PLE outer layers have improved association to ovarian and glioma cells.^18,51,52^ In this work, we found that PAA NPs displayed minimal specificity toward blood-circulating immune cells and major organs in vivo, attributed to the polymer’s bioinert nature and lack of known binding partners. By contrast, HA NPs presented a significantly higher affinity for granulocytes and monocytes, possibly due to the high expression of the HA binding partner CD44 in immune cells.^54^ Ultimately, the stealth properties achieved with PAA layering makes this polymer outer layer ideal for our Ab–LbL NP platform.

The clinical need for HSPC-targeting strategies has driven expanded efforts to identify promising targets for these cells. We sought to better understand and compare the advantages and limitations of different targeting moieties in the context of ex vivo and in vivo delivery. We rationally selected a library of HSPC-targeting Abs (anti-cKit, anti-CD45, anti-CXCR4, anti-CD90, and anti-CD105), and optimized a method to facilitate highly efficient conjugation of these Abs to PAA NPs without major changes in the physicochemical properties of the resulting Ab–PAA NPs. Functionalization with each of the Abs modified the performance of the Ab–PAA NPs in bone marrow-resident cells in vitro, both in terms of uptake and trafficking. In particular, cKit–PAA NPs demonstrated high association rates and high internalization at short time points, likely due to the rapid turnover of the receptor.^48^ Previous functionalization of LNPs with cKit has similarly increased transfection rates in HSPCs.^30,31^ In the case of anti-CD45, some studies have shown an effective HSPC targeting capacity in vivo,^25,27^ while others have not seen significant targeting or delivery improvements.^30^ Here, our work suggests the low uptake of CD45–PAA NPs may be related to the receptor’s low internalization. Other drug delivery strategies have in fact exploited CD45 to anchor nano- or microparticles onto cell surfaces to avoid internalization.^55,56^ In our hands, anti-CD90 demonstrated comparable or superior in vitro targeting properties to anti-cKit, although with a significantly different percentage of receptors expressed by bone marrow cells and different internalization patterns. These improvements could be leveraged for ex vivo cell manipulation, enabling a more effective targeted delivery to hematopoietic progenitors.

Nevertheless, in vivo HSPC-targeted delivery remains the ultimate therapeutic goal. From our findings, cKit– and CD90—PAA NPs still presented some increased targeting in vivo, while CD105–PAA NPs demonstrated a unique and remarkable in vivo targeting capacity not recapitulated in vitro. The expression of the CD105 receptor on the surface of long-term HSCs, on endothelial cells, and in the bone marrow niche, might have facilitated the localization of these NPs to the bone marrow and, hence, provided an improved targeting of HSPCs.^37,57,58^ It is worth noting that in other studies cKit–conjugated lipid nanoparticle systems showed significant improvements in HSPC transfection in vivo, as well as lung tropism not seen here.^30,31^ Variations in the NP core – in this case, a liposome rather than a lipid nanoparticle complexed with RNA – could explain these differences, as well as other factors that could vary such as Ab dose, or endpoint. Based on this premise, here we highlight the therapeutic potential of anti-CD105 for in vivo HSPC targeting, though we note that anti-CD90 and anti-cKit may perform strongly in other contexts. In parallel, we also observed higher accumulation of the most successful systems in some circulating immune cells. The heterogeneity of receptor expression across hematopoietic lineages limits the ability to achieve fully exclusive targeted delivery of NPs. Nevertheless, even a modest increase in drug uptake by targeted cells can yield substantial therapeutic benefits. Further specificity can be achieved for certain genetic cargos with tissue-specific promoters. Additionally, the net efficacy of CD105–PAA NPs could be boosted through HSPC mobilization, administration of multiple doses, or evaluation over longer time points.

Multiple clinical and preclinical interventions have leveraged the expression of various cellular markers to enhance specificity or address patient heterogeneity, using either multiple ligands or bispecific Abs.^59,60^ Building on this work, we aimed to evaluate dual-targeted Ab–PAA NPs by incorporating combinations of the three most promising targeting moieties. In vivo, none of the tested Ab combinations improved the performance of the single-targeted CD105–PAA NPs. The absence of complementary effects observed could be attributed to differences in receptor expression and internalization patterns for the different receptors, as seen in vitro. Additional factors, such as the distance between receptors, the Ab ratio, and interactions with a greater number of off-target cells, may explain these results.^59^

Notably, CD105–PAA NPs showed the highest association to healthy human CD34+ cells, suggesting the clinical translatability of this platform. This trend aligns with reports of anti-human CD105–NPs claiming good transfection rates, although the performance of this system was not directly compared to other targeting strategies.^31,41^ The variability of results among donors demonstrates that there are still limitations in finding one single-targeted strategy that can fit many patients. Personalized medicine approaches based on a characterization of the patients’ marker expression could contribute to designing more effective clinical solutions.^61^

We sought to broaden the applications of the Ab–PAA NP platform to human blood malignancies, as certain leukemia and lymphoma cells share markers with HSPC and other hematopoietic lineages.^62,63^ Similarly, targeted strategies can decrease side effects mediated by unwanted myeloid uptake, alleviating toxicity of current clinical treatments. Recent approaches have functionalized clinically-available PEG NPs with bispecific Abs targeting different common hematopoietic receptors (e.g., CD20, CD22, or CD38) to mitigate these side effects.^64,65^ However, uptake by myeloid cells was not decreased.^65^ Here, CXCR4 receptor expression was confirmed across three different human B-cell-derived cell lines —B-cell ALL, diffuse large B-cell lymphoma, and Burkitt lymphoma–,^66^ which consequently improved the uptake of CXCR4–PAA NPs. This targeting capacity, combined with their minimal interactions with murine myeloid cells and negligible accumulation in major organs, indicated that CXCR4–PAA NPs hold great potential as a selective delivery system for targeting malignancies while sparing healthy cells. Furthermore, the LbL NP platform avoids PEG derivatives in the liposomal core by leveraging the polymer layers for stability and prolonged circulation,^67^ minimizing the risk of off-target myeloid uptake caused by circulating anti-PEG Abs.^65^

This systematic study has also helped elucidate other targeting properties of LbL NPs and Ab– LbL NPs with interest beyond the scope of this work. For instance, HA NPs exhibited an increased association to T cells and to the spleen in vivo in healthy mice. Functionalization with anti-CD90 also enhanced in vivo targeting of circulating T cells, which can be associated to its expression by and internalization in peripheral T cells.^55,68^ By contrast, cKit–PAA NPs presented an enhanced targeting to B cells. Some reports suggest the presence of the cKit receptor in certain B-cell precursors.^69^

Overall, we have developed a modular Ab–LbL NP system with stealth properties capable of targeting hematopoietic progenitors in healthy and diseased states by exchanging a rationally selected targeting moiety. This platform facilitates a methodical comparison of different targeting approaches by only changing the outer surface without affecting any other parameters. Using this strategy, we identified the merits and limitations of multiple HSPC-targeting Ab candidates, suggesting that their efficacy is context-dependent to ex vivo and in vivo applications and disease states. These results emphasize the importance of systematic studies to better understand how to leverage the biology of the tissues of interest to develop optimized drug delivery systems with potential therapeutic applications.

## MATERIALS AND METHODS

### Materials

Cholesterol, 1,2-distearoyl-sn-glycero-3-phospho-(1’-rac-glycerol) sodium salt (DSPG), 1,2-distearoyl-sn-glycero-3-phosphoethanolamine (DOPE), 1,2-distearoyl-sn-glycero-3-phosphocholine (DSPC), and 1,2-distearoyl-sn-glycero-3-phosphoethanolamine-N-(Cyanine 5) (DSPE-Cy5) were purchased from Avanti (Alabama, US).

Sulfo-cyanine dyes with NHS ester were purchased from Lumiprobe (Maryland, US). Tangential flow filtration filters were purchased from Repligen (Massachusetts, US).

Poly-L-arginine hydrochloride (19 kDa), poly-L-aspartic acid (14 kDa), and poly-L-glutamic acid (15 kDa) were purchased from Alamanda Polymers (Alabama, US). Hyaluronic acid (20 kDa) was purchased from LifeCore Biomedical (Minnesota, US). Polyacrylic acid (15 kDa) was purchased from Sigma-Aldrich (St. Louis, US).

Carboxylated polystyrene nanoparticles (FluoSpheres^TM^, yellow-green) of 100 nm were purchased from Thermo Fisher Scientific (Massachusetts, US). Electron microscopy Carbon support film 200 mesh copper were obtained from Electron Microscopy Sciences (Pennsylvania, US). Goat-anti-Rat IgG (H+L0 6-nm gold nanoparticles were purchased from AURION Immuno Gold Reagents & Accessories (The Netherlands).

The antibodies rat-anti-mouse-cKit (CD117, clone 2B8), mouse-anti-mouse-CD45.2 (clone 104.2), rat-anti-mouse CD90.2 (Thy1.2, clone 30H12) and the corresponding isotypes (mouse IgG2a C1.19.4 and rat IgG2b LTF-2) were purchased from BioXCell (New Hampshire, USA). Antibody rat-anti-mouse-CXCR4 (CD184, clone 2B11), rat-anti-mouse CD105 (clone MJ7/18), mouse-anti-human CD45 (clone HI30), mouse-anti-human CXCR4 (CD184, clone 12G5), mouse-anti-human CD90 (Thy-1, clone 5E10), and mouse-anti-human CD105 (clone SN6) were purchased from Thermo Fisher Scientific (Massachusetts, US). Antibody mouse-anti-human-cKit (clone 104D2) was purchased from Peprotech (New Jersey, US).

Maleimide-PEG3-tetraciclooctane, AZDye488–tetrazine, and MethylTetrazine-amine were purchased from Click Chemistry Tools (Arizona, US). N-hydroxysuccinimide (NHS) and N-(3-(dimethylamino)-propyl)-N-ethylcarbodiimide hydrochloride (EDC) were bought from Sigma-Aldrich (St. Louis, US),

Roswell Park Memorial Institute (RPMI 1640) cell culture media and penicillin/streptomycin were purchased from Corning (New York, US), and fetal bovine serum (FBS) was purchased from Gibco (Montana, US). StemSpan SFEM II serum-free, L-glutamine, recombinant human stem cell factor (SCF), recombinant mouse and human Fms-like tyrosine kinase 3/fetal liver kinase-2 (Flt3/Flk-2) ligands, recombinant mouse and human interleukin-3 (IL-3), recombinant mouse thrombopoietin (TPO) and human CD34+ cells were purchased from Stemcell Technologies (British Columbia, Canada). Recombinant mouse stem cell factor (mSCF) and recombinant human thrombopoietin (hTPO) were obtained from Peprotech (New Jersey, US).

The 8-Chamber Lab-Tek II Chambered #1.5 German Coverglass system, wheat germ agglutinin Alexa Fluor™ 555 and 647, and Hoescht 33342 trihydrochloride trihydrate were purchased from Invitrogen, (Massachusetts, US). A 0.01% poly-L-lysine solution was purchased from Sigma-Aldrich (St. Louis, US). Hank’s balanced salt solution (HBSS) was obtained from Gibco (Montana, US), and 16% methanol-free formaldehyde from Thermo Fisher Scientific (Massachusetts, US).

### Liposome synthesis

Liposomes were prepared as previously described.^70^ Cholesterol and lipid stocks were made in only chloroform, or a chloroform : methanol : water [65 : 35 : 8 ratio] mixture, and combined in a flask at a mol ratio of 31 DSPC : 31 cholesterol : 31 DSPG : 7 DOPE or of 33 DSPC : 33 cholesterol : 33 DSPG : 0.4 DSPE-Cy5. The lipids were dried into a thin film using a BUCHI RotoVap system under heat (55 °C, water bath) until completely dry (<30 mBarr). A Branson sonicator bath was heated to 65 °C, at which point the flask with the lipid film was partially submerged in the bath. A volume of ultrapure water was added to re-suspend the lipid film to a 2 mg lipid/mL solution. The liposome solution was sonicated on and off for 3 minutes, then transferred to an Avestin LiposoFast LF-50 liposome extruder. The extruder was connected to a Cole-Parmer Polystat Heated Recirculator Bath to maintain a temperature of 65 °C. The liposomes were extruded through nucleopore membranes by passing the solution through stacked 400 and 200 nm membranes, 100 nm, and 50 nm membranes. Fluorescently labeled liposomes were prepared either with a Cy5-dye, or via NHS-coupling of Sulfo-cyanine dyes with NHS ester handles to DOPE head group amines in sodium carbonate buffer (pH 8.5), stirred at room temperature overnight. Unconjugated dye was purified away from the labeled liposomes by tangential flow filtration (TFF).

### Layer-by-Layer assembly

Nanoparticles were layered with polymers by adding an equal volume of NP core (at either 1 mg/mL or 0.5 mg/mL for HA) to the polymer solution under sonication for a few seconds.^70^ The weight equivalents of polyelectrolyte used with respect to liposome core were of 0.25-0.33 for poly-L-arginine (PLR), 1-2 for hyaluronic acid (HA), 0.5-1 for poly-L-glutamic acid (PLE), and 0.125-0.33 for polyacrylic acid (PAA). For antibody-conjugated NPs, the final layer was a 1:1 mass ratio of unmodified PAA to methyltetrazine-modified PAA. Polyelectrolyte solutions were prepared in 50 mM HEPES (pH 7.4) and 40 mM NaCl, with the exception of HA, which was prepared in 2.5 mM HEPES.^70^ The freshly layered particles were allowed to incubate at room temperature for 30 min, then purified using tangential flow filtration (TFF).

### Polymer functionalization

Carboxylated polymer poly(acrylic acid) (PAA) wa modified with a methyltetrazine handle by NHS/EDC coupling chemistry. PAA was diluted in MES buffer (0.1 N, pH 6) to a concentration of 2 mg/mL. Then, solutions of methyltetrazine amine or 3-azidopropylamine, N-hydroxysuccinimide (NHS), and N-(3-(dimethylamino)-propyl)-N-ethylcarbodiimide hydrochloride (EDC) in the same buffer were added at a concentration 6.3-, 14.3-, and 143-fold times higher than the polymer, respectively. The solution was kept under magnetic stirring at 700 rpm for 4 h at room temperature and then dialyzed against NaCl 50 mM overnight, and up to 72 h against ultrapure water to remove free reactives (Spectrum Labs, regenerated cellulose membrane MW 3.5 kDa, California, US). The final polymer solution was frozen at −80 °C and lyophilized in a Labconco FreeZone Freeze Dryer System (Labconco, Missouri, USA).

Lyophilized polymers were resuspended in D2O (Sigma-Aldrich, St. Louis, US), and analyzed by nuclear magnetic resonance (NMR). All NMR spectra were obtained using a three-channel Bruker Avance Neo spectrometer 500.34 MHz equipped with a 5 mm liquid-nitrogen cooled Prodigy broad band observe cryoprobe. Spectra were processed using MestReNova v14.2 (Mestrelab Research, Santiago de Compostela, Spain).

### Purification by tangential flow filtration

Nanoparticles were purified using a Spectrum Labs KrosFlo II filtration system (Repligen, Massachusetts, USA). Hollow fiber filter membranes with 100 or 300 kDa molecular weight cutoffs made of modified polyethylene sulfone were used to remove free dyes or polymers from the nanoparticle solution. Samples were filtered at 13 mL/min or 80 mL/min (for 13 or 16 tubing, respectively), with an ultrapure water inlet line to replace the volume of waste permeate. Nanoparticles were considered purified after collecting at least 5 volume equivalents of waste (or 10 for the antibody-conjugated ones). Then, the sample was concentrated to the desired volume.

After each purification step, nanoparticle concentration was tracked based on a calibration curve of the initial liposome solution measuring the Cy5 fluorescent reading (excitation/emission 630/670 nm). For this purpose, nanoparticle samples were diluted in a 1 : 10 ratio in dimethyl sulfoxide, and read in a Tecan M1000 microplate reader (Tecan, Männedorf, Switzerland).

### Nanoparticle characterization

The mean size and polydispersity index (PDI) of the nanoparticles were characterized by dynamic light scattering (DLS), and the zeta potential values by laser doppler anemometry, using a Malvern Zetasizer Pro (Malvern Panalytical, Malvern, UK). The measurements were done in triplicate, at 25 °C, with a red laser (landa=633nm) and a detection angle of 173°.

For cryo-electron microscopy (Cryo-EM), 3 uL of the NP sample and buffer containing solution were dropped on a lacey copper grid coated with a continuous carbon film and blotted to remove excess sample without damaging the carbon layer by Gatan Cryo Plunge III. Grid was mounted on a Gatan 626 single tilt cryo-holder equipped in the TEM column. The specimen and holder tip were cooled down by liquid-nitrogen. Imaging on a JEOL 2100 FEG microscope was done using minimum dose method that was essential to avoid sample damage under the electron beam. The microscope was operated at 200 kV and with a magnification in the ranges of 10,000–60,000 for assessing particle size and distribution. All images were recorded on a Gatan 2kx2k UltraScan CCD camera.

### Antibody modification and conjugation to nanoparticles

Full antibodies were partially reduced by using 4-molar excess (to the antibody) of tris(2-carboxyethyl)-phosphine hydrochloride (TCEP) in a final buffer solution of 20 mM phosphate buffer, 150 mM NaCl, and 10 mM EDTA (pH 7.2). The mixture was incubated for 1 h at 37 °C, and then a 40-molar excess of the maleimide-PEG3-TCO linker was added. The reaction was incubated for 1 h at room temperature, and then purified using a 7K Zeba™ Spin Column (Thermo Fisher Scientific, Massachusetts, US).

The number of linkers per antibody was approximated by conjugating an AZDye488™-tetrazine to the antibody and calculating the degree of labelling (DOL). For this, antibody-TCO was incubated with a 2-molar excess solution of the fluorophore for at least 1.5 h, and then purified with 7K Zeba™ Spin Column. Degree of labelling was quantified using a NanoDrop 1000 Spectrophotometer (Thermo Fisher Scientific, Massachusetts, US) reading the absorbance at 495 nm (AZDye488™-tetrazine ε = 83 000 M^-1^ cm^-1^), and the protein absorbance at 280 nm for IgG.

For the conjugation of the antibody to nanoparticles, a 0.5 mg/mL solution of nanoparticles was mixed with 5% volume of the modified antibody at 1 mg/mL, and reacted for 1-2 h at room temperature or overnight at 4°C. After this time, unconjugated antibody was removed via TFF (10 volumes waste collected).

Protein concentrations were determined based on the protein absorbance at 280 nm for IgG using a NanoDrop 1000 Spectrophotometer. For concentrations of proteins conjugated to the nanoparticles, the colorimetric – bicinchoninic acid (BCA) assay (Thermo Fisher, Scientific, Massachusetts, US) was used. In this study, all samples were diluted to the same final nanoparticle concentration, to prevent interference of the dye with the assay.

### Room temperature Transmission Electron Microscopy (TEM) and secondary staining of antibodies with gold nanoparticles

PS NPs were layered and conjugated to cKit antibody as described for liposomes. The purified cKit-PAA PS NPs and PAA PS NPs were analyzed by room temperature TEM. A 5 µL drop of NPs (0.25 mg/mL, by fluorescence) was added to a 200 mesh copper grid coated with a continuous carbon film and let sit for 15 min. After this, the excess of NP was removed with a paper filter. Gold NPs were diluted 1/20 in 1x PBS with 0.1% BSA, pH 7.4 and the grid was incubated in a 100 µL droplet for 2 h at room temperature. After this time, the grid was washed six times in 1x PBS 0.1% BSA pH 7.4; six times in 1x PBS, and six times in deionized water. Then, grids were transferred to a droplet of uranyl acetate (3% in water) and washed in ten droplets of deionized water. Excess water was removed with a filter paper, and then stored at room temperature until imaged. A JEOL 2100 FEG microscope was used in the same conditions as previously described.

The number of gold NPs per NP core was counted using FIJI. The number of PS NPs and of gold NPs touching (on top or next to) them were counted, and an average of Ab (= 1 gold NP) per PS NP was estimated. The average of 13 images, containing a total of 334 PS NPs was used for these calculations. Counting was done manually. Given the conjugation yield of the cKit-PAA PS NPs was 3.3 times lower than for the cKit-PAA NPs with a liposomal core reported in the manuscript, the final number of Abs per NPs was corrected accordingly: multiplied by 3.3.

### Bone marrow harvesting

Mice were euthanized following approved protocols. Mouse leg bones were isolated following stablished protocols ^71,72^. Isolated bones were kept in cold RPMI, and then, epiphyses were cut, and bones were spun down on a 70-µm strainer for 20 min at 1,000 g. After this step, cells were incubated with Ammonium-Chloride-Potassium (ACK) lysing buffer (Thermo Fisher Scientific, Massachusetts, US) for 5 min at room temperature, and then white blood cells were collected by centrifugation at 300 g for 5 min. Media was aspirated, then cells were washed with cold PBS and isolated by centrifugation at 300 g for 5 min. For final experiments with bone marrow cells, the cell pellet was resuspended in mouse HSC media, containing StemSpan SFEM II serum-free medium supplemented with 1% L-glutamine, 1% penicillin/streptomycin, 100 ng/mL mouse SCF, 100 ng/mL mouse Flt3/Flk-2 ligand, 20 ng/mL of mouse IL-3, and 50 ng/mL mouse TPO. Cells were seeded at a density of 50,000 cells per 200 µL per well of a 96 well plate.

### Human CD34+ cell experiments

Human CD34+ cells were thawed and cultured overnight in at a concentration of 10^6^ cells/mL in human HSC media: StemSpan SFEM II serum-free medium supplemented with 1% L-glutamine, 1% penicillin/streptomycin, 100 ng/mL human SCF, 100 ng/mL human Flt3/Flk-2 ligand, 20 ng/mL of human IL-3, and 50 ng/mL human TPO. The next day, cells were harvested, counted, and resuspended in human HSC media without cytokines at a density of 50,000 cells per 200 µL per well of a 96 well plate.

### Human leukemia cells

Human NALM6, clone G5, Toledo, and Raji cells were cultured according to manufacturer specifications in RPMI-1640 media supplemented with 10% FBS, 1% penicillin/streptomycin. Cells were incubated at 37 °C with an atmosphere with 100% humidity and 5% CO2. Cells were authenticated using STR profiling and tested for mycoplasma. For in vitro uptake studies, 40,000 NALM6 cells were seeded in 200 uL per well, and 40,000 Toledo or Raji cells were seeded in 100 uL per well.

### In vitro uptake studies

For flow cytometry experiments, cells were plated as previously described in a 96-well plate and let to grow overnight. Next day, a solution of the nanoparticles in ultrapure water were added, as a 10% of the well volume. Nanoparticles and cells were incubated for 0.5, 1.5, or 4 h. After this, cells were transferred to a non-treated 96-well plate, washed with PBS, stained for live/dead with Zombie Violet™ Fixable Viability Kit (BioLegend, California, US). For bone marrow cells, cells were then blocked for 10 min at room temperature (TruStain FcX™ (anti-mouse CD16/32) Antibody) and stained with an FITC anti-mouse Lineage Cocktail (BioLegend, California, US) for 20 min at 4 °C. Cells were then washed and resuspended in FACS buffer to be read in a flow cytometer BD LSR II or BD Fortessa (BD Biosciences, New Jersey, US). FlowJo™ v10 (BD Biosciences, New Jersey, US) was used to analyze the resulting data. Median fluorescence intensity (MFI) values are represented as the fold increase to the untargeted PAA NPs [MFIAb-NPs / MFIPAA NPs].

### Confocal imaging

The chambered slides were coated with a solution of 0.01% poly-L-lysine solution for 15 min, and then aspirated. Chambers were allowed to fully dry overnight at 37 °C. In parallel, mouse bone marrow cells were cultured on 96-well plates at 50,000 cells per 200 µL per well. The next day, cells were incubated with Cy5-labelled NPs for 1.5 h at 37 °C, 5% CO2. After this, cells were spun down, and washed with PBS twice. Then, cells were transferred to the poly-L-lysine coated chambered slides and let incubate for 15 min at room temperature. Cells were then fixed with 4% formaldehyde in PBS for 15 min at room temperature, and washed three times with HBSS over 15 min. Samples were stained with 10 µg/mL wheat germ agglutinin labelled with Alexa Fluor™ 555 for 10 min. Then, they were washed again with PBS three times over 15 min, stained with 1.5 µg/mL Hoechst 33342 for 2 min. Finally, samples were washed with PBS and stored at 4 °C and protected from light until imaging. Imaging was conducted in an Olympus FV1200 Laser Scanning Confocal Microscope with an inverted IX83 microscope equipped with 405, 440, 473, 559, and 635 nm lasers. Images were acquired with 100x objective. Final image analysis was done with FIJI.

### Animal husbandry

All animal experiments were approved by the MIT Committee for Animal Care (protocol 221000434). Healthy female C57BL/6J mice 6−8 weeks old were obtained from The Jackson Laboratory and housed at the Koch Institute animal facility at MIT, in groups of up to 5 mice per cage. Animals were allowed to acclimate for at least 72 h before any manipulation. Routine husbandry was provided with the assistance of the MIT Division of Comparative Medicine (DCM) staff. Animals were given ad libitum access to IsoPro RMH 3000 chow (LabDiet) and water and were maintained in 12 h light-dark cycles.

### In vivo nanoparticle dosing

Female C57BL/6J mice 6−8 weeks old were injected retro-orbitally with Cy5-labeled LbL NPs at a lipid concentration of 0.75 mg/mL; and an Ab concentration of up to 30 mg/kg (<0.2 mg/mL) in 200 uL. All NPs were dosed in sterile-filtered isotonic 5% dextrose. At the corresponding time points, blood was collected via retro-orbital bleeding (not in the eye of injection), and mice were euthanized.

### Organ harvesting and fluorescence analysis

Organs were harvested, immersed in RPMI, dried and weighed, then transferred to cold PBS. Organs were kept on ice for all intermediate periods. To determine fluorescent signal, excised organs were dried and imaged on a Xenogen IVIS Imaging System (PerkinElmer, Massachusetts). Fluorescence is reported as the total radiant efficiency (in [p/s] / [μW/cm^2^]), normalized by the values of the control organs.

### Blood analysis by flow cytometry

Blood was collected in a BD Microtainer® tube with K2 EDTA (BD Biosciences, New Jersey, US). Within 1h of the collection, tubes were centrifuged for 20 min at 1,000 g. Supernatant was removed, and remaining cells were incubated in ACK lysing buffer for 5 min, then centrifuged for 5 min at 300 g. This process was repeated until a white cell pellet withwas obtained. Cells were then resuspended in 1 mL of RPMI and left in ice until flow preparation. Within 3 h, blood cells were counted, and 10^6^ cells per mouse were plated in a 96-well plate. Cells were washed with PBS, stained for live/dead with Zombie Aqua™ Fixable Viability Kit, then blocked for 10 min at room temperature (TruStain FcX™) and stained with a cocktail containing FITC-CD45 (clone 30-F11, 103107), Pacific Blue™-B220 (clone RA3-6B2, 103230), PE-CD3 (clone 17A2, 100205), and PE/Cyanine7-CD11b (clone M1/70, 101215) in Cell Staining Buffer (BioLegend, California, US) for 25 min at 4 °C. All staining ratios used followed manufacturer recommendations. Cells were then washed twice and resuspended in Cell Staining Buffer to be analyzed in a flow cytometer BD Fortessa (BD Biosciences, New Jersey, US). Compensation was calculated with compensation beads (Thermo Fisher Scientific, Massachusetts, US). FlowJo™ v10 (BD Biosciences, New Jersey, US) was used to analyze the resulting data. Gating strategy is shown in Suppl. Fig. 1.

### Flow cytometry analysis of bone marrow cells from nanoparticle-dosed mice

Bone marrow was extracted as previously described in the “Bone marrow harvesting” section. Cells were resuspended in 4 mL of cold PBS and left in ice until flow preparation. Within 3 h, bone marrow cells were counted, and 3×10^6^ cells per mouse were plated in a 96-well plate. Cells were washed with PBS, stained for live/dead with Zombie Aqua™ Fixable Viability Kit, then blocked for 10 min at room temperature (TruStain FcX™) and stained with a cocktail containing Alexa Fluor ® 700 Lineage cocktail (clones 17A2; RB6-8C5; RA3-6B2; Ter-119; M1/70; 133313), FITC-cKit (clone ACK2, 135115), Pacific Blue^TM^-Sca-1 (clone E13-161.7, 122519), PE-CD48 (clone HM48-1, 103405), PE/Cyanine7-CD150 (clone TC15-12F12.2, 115914) in Cell Stanining Buffer (BioLegend, California, US) for 25 min at 4 °C. All staining ratios used followed manufacturer recommendations. Cells were then washed twice and resuspended in Cell Staining Buffer to be analyzed in a flow cytometer BD Fortessa (BD Biosciences, New Jersey, US). Compensation was calculated with compensation beads (Thermo Fisher Scientific, Massachusetts, US). FlowJo™ v10 (BD Biosciences, New Jersey, US) was used to analyze the resulting data. Gating strategy is shown in Suppl. Fig. 7.

### Statistical analysis

Data analysis was performed using GraphPad Prism version 9.1 (GraphPad Inc.). Statistical analysis is described in each figure caption. Data were expressed as the mean ± standard deviation (SD). p values of 0.05 or less were considered statistically significant.

## Acknowledgments

This work was supported, in part, by the Bill and Melinda Gates Foundation (grant ID INV-021791 and INV-050202). Under the grant conditions of the Foundation, a Creative Commons Attribution 4.0 Generic License has already been assigned to the Author Accepted Manuscript version that might arise from this submission. Research was also funded by the Koch Institute’s Marble Center for Cancer Nanomedicine. N.N is supported by the NSF Graduate Research Fellowship Program under Grant No. 2141064. Any opinions, findings, and conclusions or recommendations expressed in this material are those of the authors and do not necessarily reflect the views of the NSF. We would like to thank the Koch Institute’s Robert A. Swanson (1969) Biotechnology Center for technical support, specifically the Flow Cytometry Core, Microscopy Core, Nanotechnology and Nanomaterials core, High Throughput Sciences core, and the ES Cell and Transgenics Facility. Figures in this manuscript were partially created using https://www.Biorender.com. We would additionally like to thank Dr. Margaret Billingsley and Dr. Nicholas Lamson for their help in reviewing this manuscript.

## Competing interests

P.T.H. is a member of the Board of Alector Therapeutics and the Board of Sail Biomedicine, a Flagship company, and a former member of the Scientific Advisory Board of Moderna Therapeutics and the Board of LayerBio. All other authors report no competing interests.

## Notes

### Summary of Updates

This version of the manuscript includes updates in the results and discussion sections.

## References

(1) Cavazzana, M.; Bushman, F. D.; Miccio, A.; André-Schmutz, I.; Six, E. Gene Therapy Targeting Haematopoietic Stem Cells for Inherited Diseases: Progress and Challenges. Nat. Rev. Drug Discov. 2019, 18 (6), 447–462. 10.1038/s41573-019-0020-9.

(2) Ferrari, G.; Thrasher, A. J.; Aiuti, A. Gene Therapy Using Haematopoietic Stem and Progenitor Cells. Nat. Rev. Genet. 2021, 22 (4), 216–234. 10.1038/s41576-020-00298-5.

(3) Anthony, B. A.; Link, D. C. Regulation of Hematopoietic Stem Cells by Bone Marrow Stromal Cells. Trends Immunol. 2014, 35 (1), 32–37. 10.1016/j.it.2013.10.002.

(4) Jenq, R. R.; Van Den Brink, M. R. M. Allogeneic Haematopoietic Stem Cell Transplantation: Individualized Stem Cell and Immune Therapy of Cancer. Nat. Rev. Cancer 2010, 10 (3), 213–221. 10.1038/nrc2804.

(5) Drysdale, C. M.; Nassehi, T.; Gamer, J.; Yapundich, M.; Tisdale, J. F.; Uchida, N. Hematopoietic Stem Cell-Targeted Gene-Addition and Gene-Editing Strategies for β-Hemoglobinopathies. Cell Stem Cell 2021, 28 (2), 191–208. 10.1016/j.stem.2021.01.001.

(6) Morgan, R. A.; Gray, D.; Lomova, A.; Kohn, D. B. Hematopoietic Stem Cell Gene Therapy: Progress and Lessons Learned. Cell Stem Cell 2017, 21 (5), 574–590. 10.1016/j.stem.2017.10.010.

(7) High-Dose AAV Gene Therapy Deaths. Nat. Biotechnol. 2020, 38 (8), 910–910. 10.1038/s41587-020-0642-9.

(8) Guo, J.; Luan, X.; Cong, Z.; Sun, Y.; Wang, L.; McKenna, S. L.; Cahill, M. R.; O’Driscoll, C. M. The Potential for Clinical Translation of Antibody-Targeted Nanoparticles in the Treatment of Acute Myeloid Leukaemia. J. Controlled Release 2018, 286, 154–166. 10.1016/j.jconrel.2018.07.024.

(9) Li, J.; Wang, Q.; Han, Y.; Jiang, L.; Lu, S.; Wang, B.; Qian, W.; Zhu, M.; Huang, H.; Qian, P. Development and Application of Nanomaterials, >Nanotechnology and Nanomedicine for Treating Hematological Malignancies. J. Hematol. Oncol. 2023, 16 (1), 65. 10.1186/s13045-023-01460-2.

(10) Lieber, A.; Kiem, H.-P. Prospects and Challenges of in Vivo Hematopoietic Stem Cell Genome Editing for Hemoglobinopathies. Mol. Ther. 2023, 31 (10), 2823–2825. 10.1016/j.ymthe.2023.09.006.

(11) Murugesan, R.; Karuppusamy, K. V.; Marepally, S.; Thangavel, S. Current Approaches and Potential Challenges in the Delivery of Gene Editing Cargos into Hematopoietic Stem and Progenitor Cells. Front. Genome Ed. 2023, 5, 1148693. 10.3389/fgeed.2023.1148693.

(12) Dacoba, T. G.; Anthiya, S.; Berrecoso, G.; Fernández-Mariño, I.; Fernández-Varela, C.; Crecente-Campo, J.; Teijeiro-Osorio, D.; Torres Andón, F.; Alonso, M. J. Nano-Oncologicals: A Tortoise Trail Reaching New Avenues. Adv. Funct. Mater. 2021, 31 (44), 2009860. 10.1002/adfm.202009860.

(13) Samaridou, E.; Heyes, J.; Lutwyche, P. Lipid Nanoparticles for Nucleic Acid Delivery: Current Perspectives. Adv. Drug Deliv. Rev. 2020, 154–155, 37–63. 10.1016/j.addr.2020.06.002.

(14) Zhang, X.; Liang, T.; Ma, Q. Layer-by-Layer Assembled Nano-Drug Delivery Systems for Cancer Treatment. Drug Deliv. 2021, 28 (1), 655–669. 10.1080/10717544.2021.1905748.

(15) Boehnke, N.; Hammond, P. T. Power in Numbers: Harnessing Combinatorial and Integrated Screens to Advance Nanomedicine. JACS Au 2022, 2 (1), 12–21. 10.1021/jacsau.1c00313.

(16) Deng, Z. J.; Morton, S. W.; Ben-Akiva, E.; Dreaden, E. C.; Shopsowitz, K. E.; Hammond, P. T. Layer-by-Layer Nanoparticles for Systemic Codelivery of an Anticancer Drug and siRNA for Potential Triple-Negative Breast Cancer Treatment. ACS Nano 2013, 7 (11), 9571–9584. 10.1021/nn4047925.

(17) Gu, L.; Deng, Z. J.; Roy, S.; Hammond, P. T. A Combination RNAi-Chemotherapy Layer-by-Layer Nanoparticle for Systemic Targeting of KRAS/P53 with Cisplatin to Treat Non–Small Cell Lung Cancer. Clin. Cancer Res. 2017, 23 (23), 7312–7323. 10.1158/1078-0432.CCR-16-2186.

(18) Barberio, A. E.; Smith, S. G.; Correa, S.; Nguyen, C.; Nhan, B.; Melo, M.; Tokatlian, T.; Suh, H.; Irvine, D. J.; Hammond, P. T. Cancer Cell Coating Nanoparticles for Optimal Tumor-Specific Cytokine Delivery. ACS Nano 2020, 14 (9), 11238–11253. 10.1021/acsnano.0c03109.

(19) Morton, S. W.; Poon, Z.; Hammond, P. T. The Architecture and Biological Performance of Drug-Loaded LbL Nanoparticles. Biomaterials 2013, 34 (21), 5328–5335. 10.1016/j.biomaterials.2013.03.059.

(20) Nabar, N.; Dacoba, T. G.; Covarrubias, G.; Romero-Cruz, D.; Hammond, P. T. Electrostatic Adsorption of Polyanions onto Lipid Nanoparticles Controls Uptake, Trafficking, and Transfection of RNA and DNA Therapies. Proc. Natl. Acad. Sci. 2024, 121 (11), e2307809121.

(21) Boehnke, N.; Dolph, K. J.; Juarez, V. M.; Lanoha, J. M.; Hammond, P. T. Electrostatic Conjugation of Nanoparticle Surfaces with Functional Peptide Motifs. Bioconjug. Chem. 2020, 31 (9), 2211–2219. 10.1021/acs.bioconjchem.0c00384.

(22) Boehnke, N.; Correa, S.; Hao, L.; Wang, W.; Straehla, J. P.; Bhatia, S. N.; Hammond, P. T. Theranostic Layer-by-Layer Nanoparticles for Simultaneous Tumor Detection and Gene Silencing. Angew. Chem. Int. Ed. 2020, 59 (7), 2776–2783. 10.1002/anie.201911762.

(23) Straehla, J. P.; Hajal, C.; Safford, H. C.; Offeddu, G. S.; Boehnke, N.; Dacoba, T. G.; Wyckoff, J.; Kamm, R. D.; Hammond, P. T. A Predictive Microfluidic Model of Human Glioblastoma to Assess Trafficking of Blood–Brain Barrier-Penetrant Nanoparticles. Proc. Natl. Acad. Sci. 2022, 119 (23), e2118697119. 10.1073/pnas.2118697119.

(24) Choi, K. Y.; Correa, S.; Min, J.; Li, J.; Roy, S.; Laccetti, K. H.; Dreaden, E.; Kong, S.; Heo, R.; Roh, Y. H.; Lawson, E. C.; Palmer, P. A.; Hammond, P. T. Binary Targeting of siRNA to Hematologic Cancer Cells In Vivo Using Layer-by-Layer Nanoparticles. Adv. Funct. Mater. 2019, 29 (20), 1900018. 10.1002/adfm.201900018.

(25) Palchaudhuri, R.; Saez, B.; Hoggatt, J.; Schajnovitz, A.; Sykes, D. B.; Tate, T. A.; Czechowicz, A.; Kfoury, Y.; Ruchika, F.; Rossi, D. J.; Verdine, G. L.; Mansour, M. K.; Scadden, D. T. Non-Genotoxic Conditioning for Hematopoietic Stem Cell Transplantation Using a Hematopoietic-Cell-Specific Internalizing Immunotoxin. Nat. Biotechnol. 2016, 34 (7), 738–745. 10.1038/nbt.3584.

(26) Hermiston, M. L.; Xu, Z.; Weiss, A. CD45: A Critical Regulator of Signaling Thresholds in Immune Cells. Annu. Rev. Immunol. 2003, 21 (1), 107–137. 10.1146/annurev.immunol.21.120601.140946.

(27) Palanki, R.; Riley, J. S.; Bose, S. K.; Luks, V.; Dave, A.; Kus, N.; White, B. M.; Ricciardi, A. S.; Swingle, K. L.; Xue, L.; Sung, D.; Thatte, A. S.; Safford, H. C.; Chaluvadi, V. S.; Carpenter, M.; Han, E. L.; Maganti, R.; Hamilton, A. G.; Mrksich, K.; Billingsley, M. B.; Zoltick, P. W.; Alameh, M.-G.; Weissman, D.; Mitchell, M. J.; Peranteau, W. H. In Utero Delivery of Targeted Ionizable Lipid Nanoparticles Facilitates in Vivo Gene Editing of Hematopoietic Stem Cells. Proc. Natl. Acad. Sci. 2024, 121 (32), e2400783121. 10.1073/pnas.2400783121.

(28) Edling, C. E.; Hallberg, B. C-Kit—A Hematopoietic Cell Essential Receptor Tyrosine Kinase. Int. J. Biochem. Cell Biol. 2007, 39 (11), 1995–1998. 10.1016/j.biocel.2006.12.005.

(29) Lennartsson, J.; Rönnstrand, L. Stem Cell Factor Receptor/c-Kit: From Basic Science to Clinical Implications. Physiol. Rev. 2012, 92 (4), 1619–1649. 10.1152/physrev.00046.2011.

(30) Shi, D.; Toyonaga, S.; Anderson, D. G. *In Vivo* RNA Delivery to Hematopoietic Stem and Progenitor Cells *via* Targeted Lipid Nanoparticles. Nano Lett. 2023, 23 (7), 2938–2944. 10.1021/acs.nanolett.3c00304.

(31) Breda, L.; Papp, T. E.; Triebwasser, M. P.; Yadegari, A.; Fedorky, M. T.; Tanaka, N.; Abdulmalik, O.; Pavani, G.; Wang, Y.; Grupp, S. A.; Chou, S. T.; Ni, H.; Mui, B. L.; Tam, Y. K.; Weissman, D.; Rivella, S.; Parhiz, H. In Vivo Hematopoietic Stem Cell Modification by mRNA Delivery. Science 2023, 381 (6656), 436–443. 10.1126/science.ade6967.

(32) Philippidis, A. Magenta Halts Development, Pursues Strategic Alternatives After Patient Death. Hum. Gene Ther. 2023, 34 (5–6), 177–179. 10.1089/hum.2023.29236.bfs.

(33) Möhle, R.; Bautz, F.; Rafii, S.; Moore, M. A.; Brugger, W.; Kanz, L. The Chemokine Receptor CXCR-4 Is Expressed on CD34+ Hematopoietic Progenitors and Leukemic Cells and Mediates Transendothelial Migration Induced by Stromal Cell-Derived Factor-1. Blood 1998, 91 (12), 4523–4530.

(34) Singh, P.; Mohammad, K. S.; Pelus, L. M. CXCR4 Expression in the Bone Marrow Microenvironment Is Required for Hematopoietic Stem and Progenitor Cell Maintenance and Early Hematopoietic Regeneration after Myeloablation. Stem Cells 2020, 38 (7), 849–859. 10.1002/stem.3174.

(35) Jørgensen, A. S.; Daugvilaite, V.; De Filippo, K.; Berg, C.; Mavri, M.; Benned-Jensen, T.; Juzenaite, G.; Hjortø, G.; Rankin, S.; Våbenø, J.; Rosenkilde, M. M. Biased Action of the CXCR4-Targeting Drug Plerixafor Is Essential for Its Superior Hematopoietic Stem Cell Mobilization. *Commun*. Biol. 2021, 4 (1), 569. 10.1038/s42003-021-02070-9.

(36) Craig, W.; Kay, R.; Cutler, R. L.; Lansdorp, P. M. Expression of Thy-1 on Human Hematopoietic Progenitor Cells. J. Exp. Med. 1993, 177 (5), 1331–1342. 10.1084/jem.177.5.1331.

(37) Kays, S.-K.; Kaufmann, K. B.; Abel, T.; Brendel, C.; Bonig, H.; Grez, M.; Buchholz, C. J.; Kneissl, S. CD105 Is a Surface Marker for Receptor-Targeted Gene Transfer into Human Long-Term Repopulating Hematopoietic Stem Cells. Stem Cells Dev. 2015, 24 (6), 714–723. 10.1089/scd.2014.0455.

(38) Rokhlin, O. W.; Cohen, M. B.; Kubagawa, H.; Letarte, M.; Cooper, M. D. Differential Expression of Endoglin on Fetal and Adult Hematopoietic Cells in Human Bone Marrow. J. Immunol. 1995, 154 (9), 4456–4465. 10.4049/jimmunol.154.9.4456.

(39) Berckmueller, K.; Thomas, J.; Taha, E. A.; Choo, S.; Madhu, R.; Kanestrom, G.; Rupert, P. B.; Strong, R.; Kiem, H.-P.; Radtke, S. CD90-Targeted Lentiviral Vectors for HSC Gene Therapy. Mol. Ther. 2023, 31 (10), 2901–2913. 10.1016/j.ymthe.2023.08.003.

(40) Parhiz, H.; Weissman, D.; Kiem, H.-P.; Radtke, S.; Peterson, C. CD-90 Targeted Lipid Nanoparticles. US 20240238438A1, July 18, 2024. https://patents.google.com/patent/US20240238438A1/en.

(41) Moffett, H. F.; Coon, M. E.; Radtke, S.; Stephan, S. B.; McKnight, L.; Lambert, A.; Stoddard, B. L.; Kiem, H. P.; Stephan, M. T. Hit-and-Run Programming of Therapeutic Cytoreagents Using mRNA Nanocarriers. Nat. Commun. 2017, 8 (1), 389. 10.1038/s41467-017-00505-8.

(42) Kelly, P. M.; Åberg, C.; Polo, E.; O’Connell, A.; Cookman, J.; Fallon, J.; Krpetić, Ž.; Dawson, K. A. Mapping Protein Binding Sites on the Biomolecular Corona of Nanoparticles. Nat. Nanotechnol. 2015, 10 (5), 472–479. 10.1038/nnano.2015.47.

(43) Herda, L. M.; Hristov, D. R.; Lo Giudice, M. C.; Polo, E.; Dawson, K. A. Mapping of Molecular Structure of the Nanoscale Surface in Bionanoparticles. J. Am. Chem. Soc. 2017, 139 (1), 111–114. 10.1021/jacs.6b12297.

(44) Challen, G. A.; Pietras, E. M.; Wallscheid, N. C.; Signer, R. A. J. Simplified Murine Multipotent Progenitor Isolation Scheme: Establishing a Consensus Approach for Multipotent Progenitor Identification. Exp. Hematol. 2021, 104, 55–63. 10.1016/j.exphem.2021.09.007.

(45) Scala, S.; Aiuti, A. In Vivo Dynamics of Human Hematopoietic Stem Cells: Novel Concepts and Future Directions. Blood Adv. 2019, 3 (12), 1916–1924. 10.1182/bloodadvances.2019000039.

(46) Bahal, R.; Ali McNeer, N.; Quijano, E.; Liu, Y.; Sulkowski, P.; Turchick, A.; Lu, Y.-C.; Bhunia, D. C.; Manna, A.; Greiner, D. L.; Brehm, M. A.; Cheng, C. J.; López-Giráldez, F.; Ricciardi, A.; Beloor, J.; Krause, D. S.; Kumar, P.; Gallagher, P. G.; Braddock, D. T.; Mark Saltzman, W.; Ly, D. H.; Glazer, P. M. In Vivo Correction of Anaemia in β-Thalassemic Mice by γPNA-Mediated Gene Editing with Nanoparticle Delivery. Nat. Commun. 2016, 7 (1), 13304. 10.1038/ncomms13304.

(47) Ricciardi, A. S.; Bahal, R.; Farrelly, J. S.; Quijano, E.; Bianchi, A. H.; Luks, V. L.; Putman, R.; López-Giráldez, F.; Coşkun, S.; Song, E.; Liu, Y.; Hsieh, W.-C.; Ly, D. H.; Stitelman, D. H.; Glazer, P. M.; Saltzman, W. M. In Utero Nanoparticle Delivery for Site-Specific Genome Editing. Nat. Commun. 2018, 9 (1), 2481. 10.1038/s41467-018-04894-2.

(48) Nagamachi, A.; Kikuchi, J.; Kanai, A.; Furukawa, Y.; Inaba, T. Kinetics of Cytokine Receptor Internalization under Steady-State Conditions Affects Growth of Neighboring Blood Cells. Haematologica 2020, 105 (7), e325–e327. 10.3324/haematol.2019.232959.

(49) Cruz, L. J.; Rezaei, S.; Grosveld, F.; Philipsen, S.; Eich, C. Nanoparticles Targeting Hematopoietic Stem and Progenitor Cells: Multimodal Carriers for the Treatment of Hematological Diseases. Front. Genome Ed. 2022, 4, 1030285. 10.3389/fgeed.2022.1030285.

(50) De Lázaro, I.; Mooney, D. J. Obstacles and Opportunities in a Forward Vision for Cancer Nanomedicine. Nat. Mater. 2021, 20 (11), 1469–1479. 10.1038/s41563-021-01047-7.

(51) Correa, S.; Boehnke, N.; Barberio, A. E.; Deiss-Yehiely, E.; Shi, A.; Oberlton, B.; Smith, S. G.; Zervantonakis, I.; Dreaden, E. C.; Hammond, P. T. Tuning Nanoparticle Interactions with Ovarian Cancer through Layer-by-Layer Modification of Surface Chemistry. ACS Nano 2020, 14 (2), 2224–2237. 10.1021/acsnano.9b09213.

(52) Pickering, A. J.; Lamson, N. G.; Marand, M. H.; Hwang, W.; Straehla, J. P.; Hammond, P. T. Layer-by-Layer Polymer Functionalization Improves Nanoparticle Penetration and Glioblastoma Targeting in the Brain. ACS Nano 2023, 17 (23), 24154–24169. 10.1021/acsnano.3c09273.

(53) Lamson, N. G.; Pickering, A. J.; Wyckoff, J.; Ganesh, P.; Calle, E. A.; Straehla, J. P.; Hammond, P. T. Trafficking through the Blood–Brain Barrier Is Directed by Core and Outer Surface Components of Layer-by-layer Nanoparticles. Bioeng. Transl. Med. 2023, e10636. 10.1002/btm2.10636.

(54) Lee-Sayer, S. S. M.; Dong, Y.; Arif, A. A.; Olsson, M.; Brown, K. L.; Johnson, P. The Where, When, How, and Why of Hyaluronan Binding by Immune Cells. Front. Immunol. 2015, 6. 10.3389/fimmu.2015.00150.

(55) Tang, L.; Zheng, Y.; Melo, M. B.; Mabardi, L.; Castaño, A. P.; Xie, Y.-Q.; Li, N.; Kudchodkar, S. B.; Wong, H. C.; Jeng, E. K.; Maus, M. V.; Irvine, D. J. Enhancing T Cell Therapy through TCR-Signaling-Responsive Nanoparticle Drug Delivery. Nat. Biotechnol. 2018, 36 (8), 707–716. 10.1038/nbt.4181.

(56) Kapate, N.; Dunne, M.; Kumbhojkar, N.; Prakash, S.; Wang, L. L.-W.; Graveline, A.; Park, K. S.; Chandran Suja, V.; Goyal, J.; Clegg, J. R.; Mitragotri, S. A Backpack-Based Myeloid Cell Therapy for Multiple Sclerosis. Proc. Natl. Acad. Sci. 2023, 120 (17), e2221535120. 10.1073/pnas.2221535120.

(57) Chen, C.-Z.; Li, M.; De Graaf, D.; Monti, S.; Göttgens, B.; Sanchez, M.-J.; Lander, E. S.; Golub, T. R.; Green, A. R.; Lodish, H. F. Identification of Endoglin as a Functional Marker That Defines Long-Term Repopulating Hematopoietic Stem Cells. Proc. Natl. Acad. Sci. 2002, 99 (24), 15468–15473. 10.1073/pnas.202614899.

(58) Kenswil, K. J. G.; Pisterzi, P.; Sánchez-Duffhues, G.; Van Dijk, C.; Lolli, A.; Knuth, C.; Vanchin, B.; Jaramillo, A. C.; Hoogenboezem, R. M.; Sanders, M. A.; Feyen, J.; Cupedo, T.; Costa, I. G.; Li, R.; Bindels, E. M. J.; Lodder, K.; Blom, B.; Bos, P. K.; Goumans, M.-J.; Ten Dijke, P.; Farrell, E.; Krenning, G.; Raaijmakers, M. H. G. P. Endothelium-Derived Stromal Cells Contribute to Hematopoietic Bone Marrow Niche Formation. Cell Stem Cell 2021, 28 (4), 653–670.e11. 10.1016/j.stem.2021.01.006.

(59) Guo, P.; Yang, J.; Liu, D.; Huang, L.; Fell, G.; Huang, J.; Moses, M. A.; Auguste, D. T. Dual Complementary Liposomes Inhibit Triple-Negative Breast Tumor Progression and Metastasis. Sci. Adv. 2019, 5 (3), eaav5010. 10.1126/sciadv.aav5010.

(60) Brinkmann, U.; Kontermann, R. E. Bispecific Antibodies. Science 2021, 372 (6545), 916–917. 10.1126/science.abg1209.

(61) Van Der Meel, R.; Sulheim, E.; Shi, Y.; Kiessling, F.; Mulder, W. J. M.; Lammers, T. Smart Cancer Nanomedicine. Nat. Nanotechnol. 2019, 14 (11), 1007–1017. 10.1038/s41565-019-0567-y.

(62) Riether, C.; Schürch, C. M.; Ochsenbein, A. F. Regulation of Hematopoietic and Leukemic Stem Cells by the Immune System. Cell Death Differ. 2015, 22 (2), 187–198. 10.1038/cdd.2014.89.

(63) Kahle, X. U.; Montes De Jesus, F. M.; Glaudemans, A. W. J. M.; Lub-de Hooge, M. N.; Jorritsma-Smit, A.; Plattel, W. J.; Van Meerten, T.; Diepstra, A.; Van Den Berg, A.; Kwee, T. C.; Noordzij, W.; De Vries, E. G. E.; Nijland, M. Molecular Imaging in Lymphoma beyond 18F-FDG-PET: Understanding the Biology and Its Implications for Diagnostics and Therapy. Lancet Haematol. 2020, 7 (6), e479–e489. 10.1016/S2352-3026(20)30065-X.

(64) Moles, E.; Howard, C. B.; Huda, P.; Karsa, M.; McCalmont, H.; Kimpton, K.; Duly, A.; Chen, Y.; Huang, Y.; Tursky, M. L.; Ma, D.; Bustamante, S.; Pickford, R.; Connerty, P.; Omari, S.; Jolly, C. J.; Joshi, S.; Shen, S.; Pimanda, J. E.; Dolnikov, A.; Cheung, L. C.; Kotecha, R. S.; Norris, M. D.; Haber, M.; De Bock, C. E.; Somers, K.; Lock, R. B.; Thurecht, K. J.; Kavallaris, M. Delivery of PEGylated Liposomal Doxorubicin by Bispecific Antibodies Improves Treatment in Models of High-Risk Childhood Leukemia. Sci. Transl. Med. 2023, 15 (696), eabm1262. 10.1126/scitranslmed.abm1262.

(65) Ju, Y.; Li, S.; Tan, A. E. Q.; Pilkington, E. H.; Brannon, P. T.; Plebanski, M.; Cui, J.; Caruso, F.; Thurecht, K. J.; Tam, C.; Kent, S. J. Patient-Specific Nanoparticle Targeting in Human Leukemia Blood. ACS Nano 2024, 18 (42), 29021–29035. 10.1021/acsnano.4c09919.

(66) Su, L.; Hu, Z.; Yang, Y. Role of CXCR4 in the Progression and Therapy of Acute Leukaemia. Cell Prolif. 2021, 54 (7), e13076. 10.1111/cpr.13076.

(67) Poon, Z.; Lee, J. B.; Morton, S. W.; Hammond, P. T. Controlling in Vivo Stability and Biodistribution in Electrostatically Assembled Nanoparticles for Systemic Delivery. Nano Lett. 2011, 11 (5), 2096–2103. 10.1021/nl200636r.

(68) Haeryfar, S. M. M.; Hoskin, D. W. Thy-1: More than a Mouse Pan-T Cell Marker1. J. Immunol. 2004, 173 (6), 3581–3588. 10.4049/jimmunol.173.6.3581.

(69) Balciunaite, G.; Ceredig, R.; Massa, S.; Rolink, A. G. A B220+ CD117+ CD19± Hematopoietic Progenitor with Potent Lymphoid and Myeloid Developmental Potential. Eur. J. Immunol. 2005, 35 (7), 2019–2030. 10.1002/eji.200526318.

(70) Correa, S.; Boehnke, N.; Deiss-Yehiely, E.; Hammond, P. T. Solution Conditions Tune and Optimize Loading of Therapeutic Polyelectrolytes into Layer-by-Layer Functionalized Liposomes. ACS Nano 2019, 13 (5), 5623–5634. 10.1021/acsnano.9b00792.

(71) Liu, X.; Quan, N. Immune Cell Isolation from Mouse Femur Bone Marrow. BIO-Protoc. 2015, 5 (20). 10.21769/BioProtoc.1631.

(72) Amend, S. R.; Valkenburg, K. C.; Pienta, K. J. Murine Hind Limb Long Bone Dissection and Bone Marrow Isolation. J. Vis. Exp. 2016, No. 110, 53936. 10.3791/53936.

