## Supporting Information for "A Modular Layer-by-Layer Nanoparticle Platform for Hematopoietic Progenitor and Stem Cell Targeting"

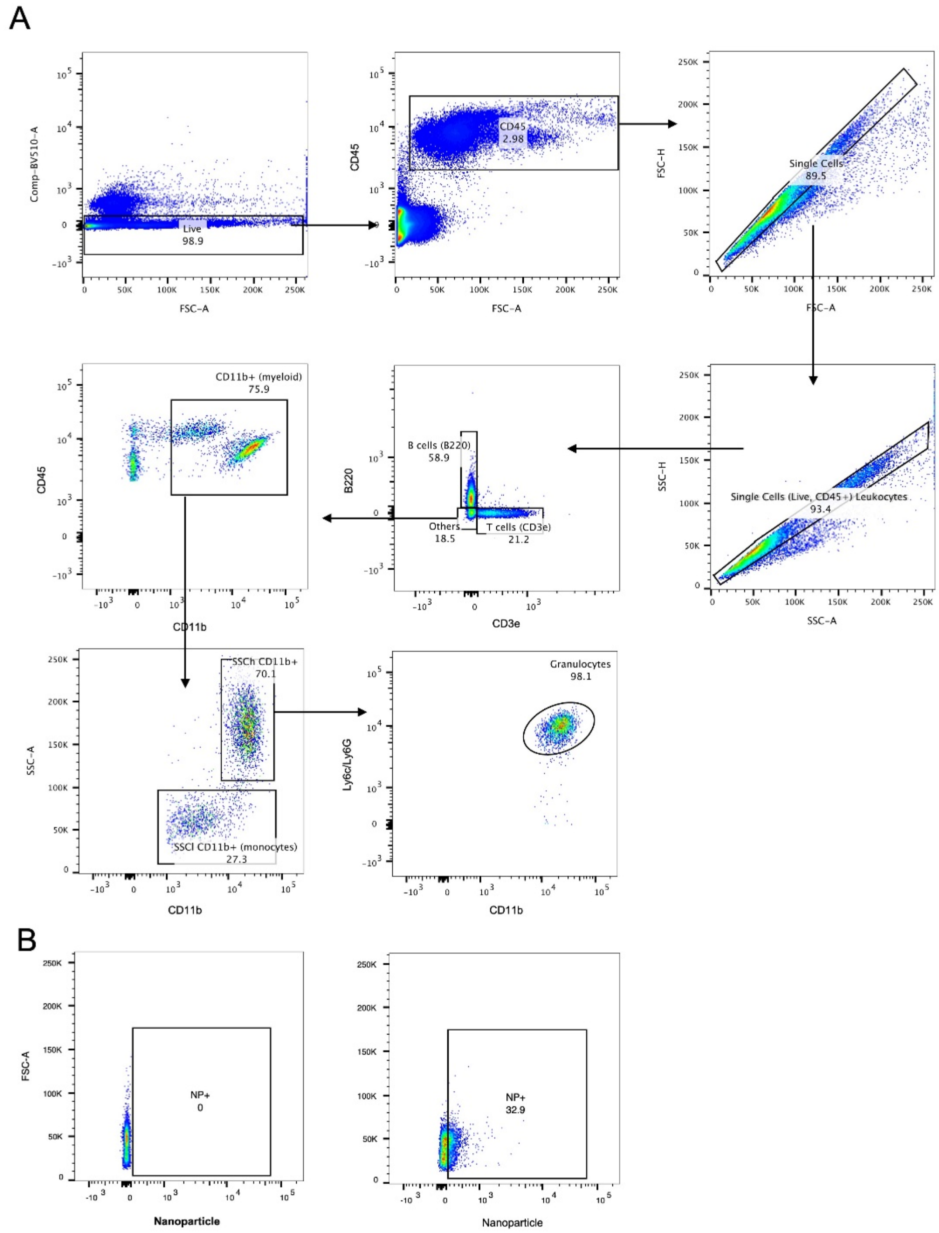


**Supporting Figure 1. Flow gating strategy used for the analysis of blood cells.** (A) Gating strategy based on different surface marker expression. (B) Representative gating for a control sample (no nanoparticle signal, left) and a sample with nanoparticle signal (right).

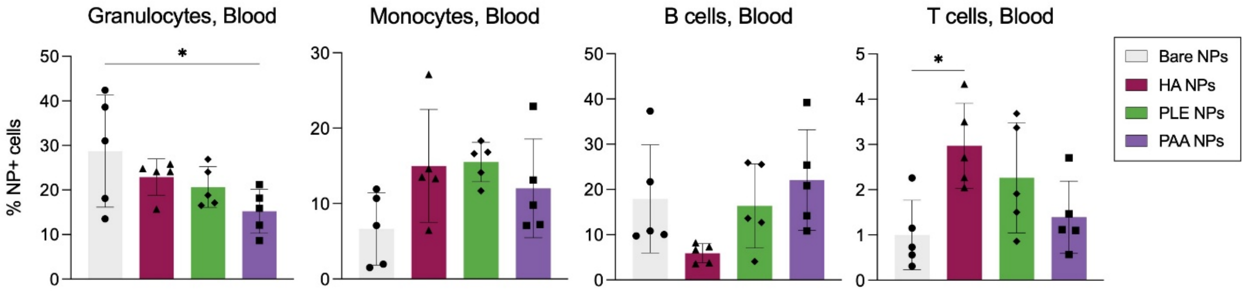


**Supporting Figure 2. LbL NP positive cells in blood cells for the different outer layers.** FACS gating strategy identifies the different cell subtypes first, and then the NP-positive populations. Percentage of cells that have taken up LbL NPs in granulocytes, monocytes, B cells, and T cells. Bars represent the mean, and each dot represents one biological replicate (n=5 per group). Statistical analysis is a one-way ANOVA with Tukey correction (*p<0.05).

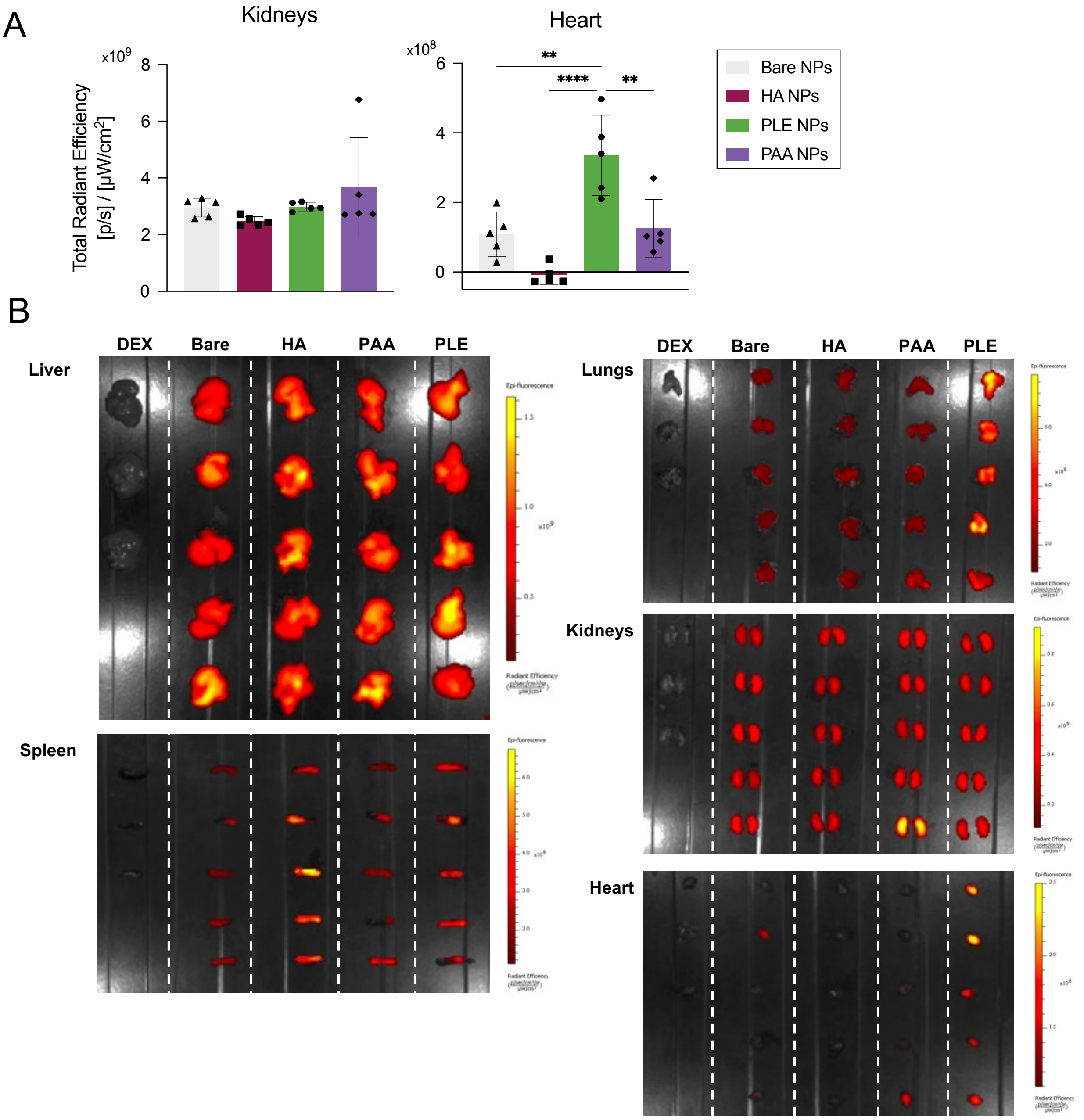


**Supporting Figure 3.** (A) Radiant efficiency measurements obtained via IVIS of the fluorophore (Cy5)-tagged Ab–PAA NPs in kidneys and heart. Bars represent the mean, and each dot represents one biological replicate (n=5 animals per group). Statistical analysis is a one-way ANOVA with Tukey correction (*p,0.05, **p<0.01, ****p<0.001). (B) IVIS images of the organs harvested 1.5 h after intravenous injection of the Ab–PAA NPs, in comparison to the 5% dextrose control (DEX) and bare liposomes (bare).

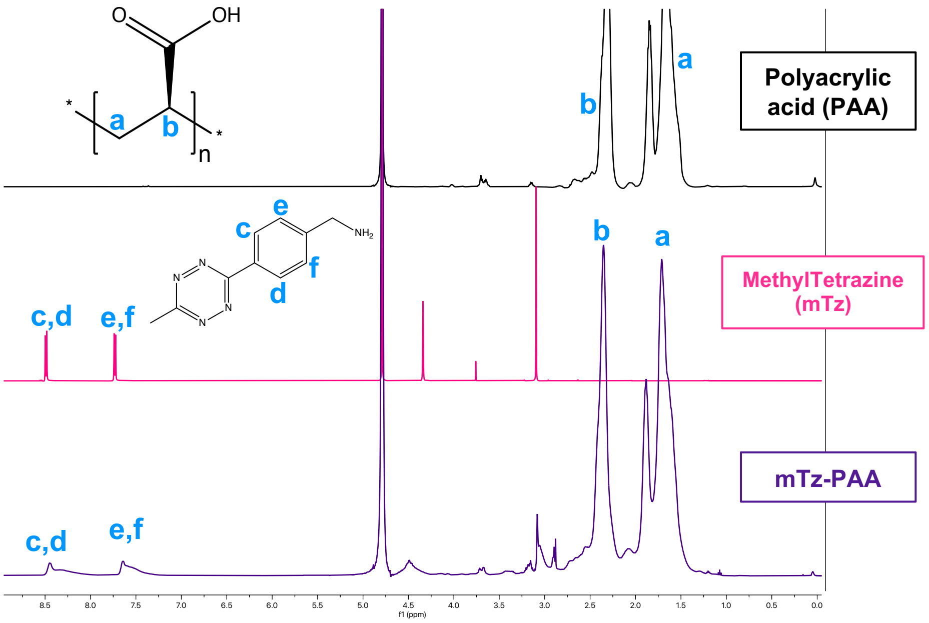


**Supporting Figure 4. Proton Nuclear Magnetic Resonance ($^1$H NMR) of polyacrylic acid (PAA), methyltetrazine amine (mTz), and the reaction conjugate methyltetrazine–polyacrylic acid (mTz-PAA).**

Polyacrylic acid (500 MHz, $H_2O$-d2): δ ppm 1.45-1.95 (d, 2H), 2.30 (s, 1H).

Methyltetrazine amine (500 MHz, $H_2O$-d2): δ ppm 7.72, 7.74 (d, 2H), 8.48, 8.50 (d, 2H),

Methyltetrazine-polyacrylic acid (500 MHz, $H_2O$-d2): δ ppm 1.36-2.00 (d, 2H), 2.35 (s, 1H), 7.28-7.68 (d, 2H), 8.00-8.50 (d, 2H).

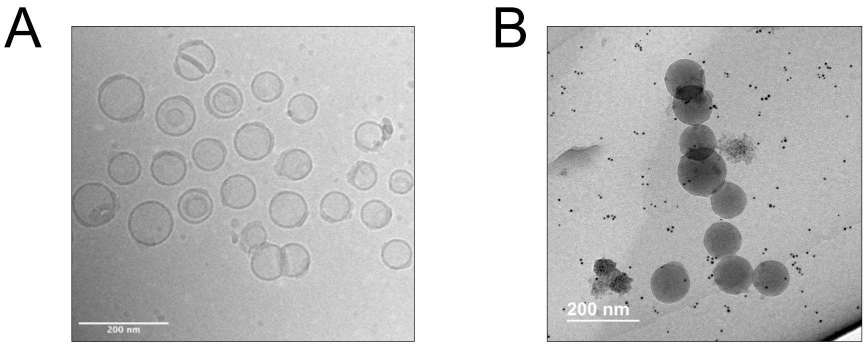


**Supporting Figure 5.** (A) Cryo transmission electron microscopy images of PAA NPs (with a liposomal core) and (B) Transmission electron microscopy images of PAA NPs (with a 100-nm polystyrene core) stained with 6-nm gold NP functionalized with an anti-rat(H&L) antibody. Scale bars represent 200 nm.

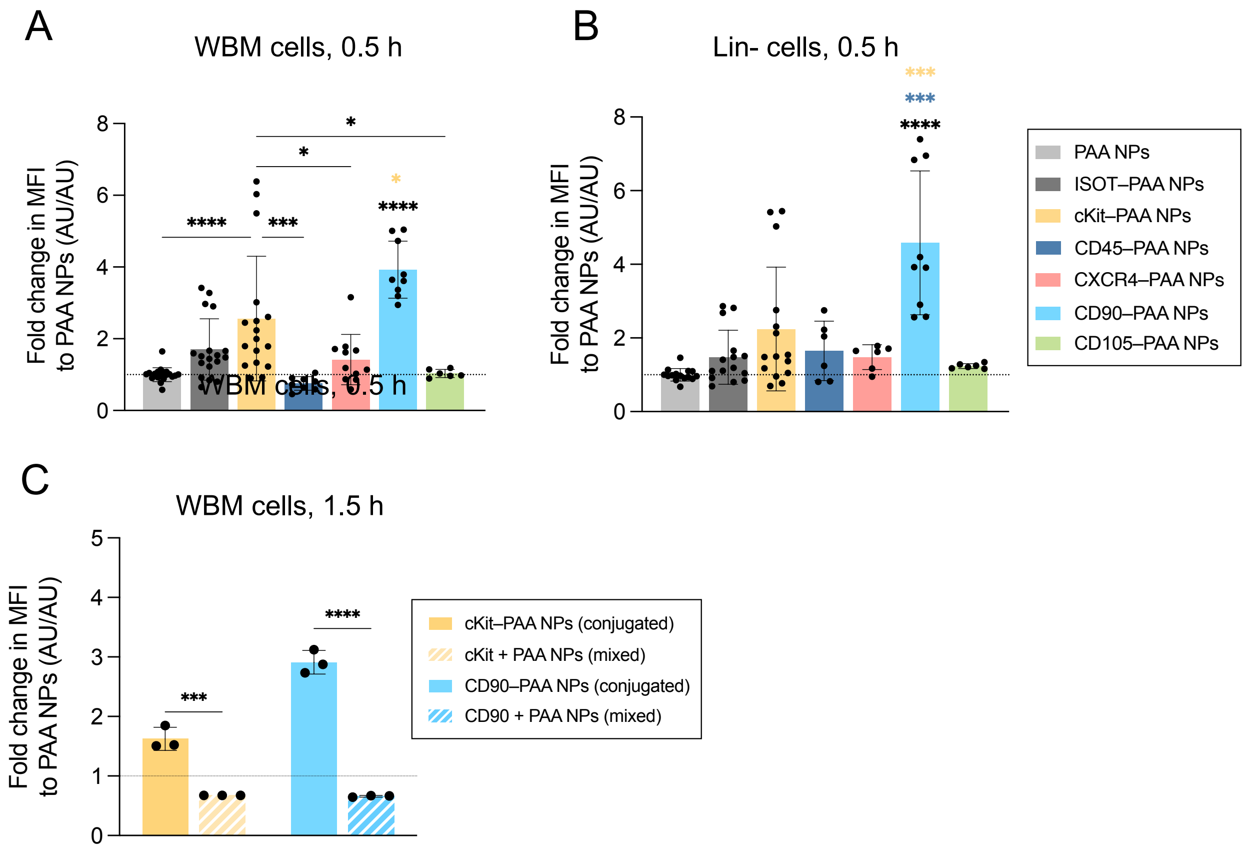


**Supporting Figure 6. Median fluorescence intensity (MFI) values after 0.5 h of incubation** with (A) WBM cells, or (B) Lin- cells, normalized to the PAA NP, for all the Ab–PAA NPs. (C) Median fluorescence intensity (MFI) values after 1.5 h of incubation with cKit– and CD90–PAA NPs in WBM cells, normalized to the PAA NP, with conjugated Abs (full bars) and mixed Abs and PAA NPs (dashed bars). Statistical analysis is a one-way ANOVA with Tukey correction (*$p<0.05$, ***$p<0.005$, ****$p<0.001$). Bars represent mean, and each dot is one technical replicate (≥2 biological replicates for A-B, 1 biological replicate for C, each with 3 technical replicates).

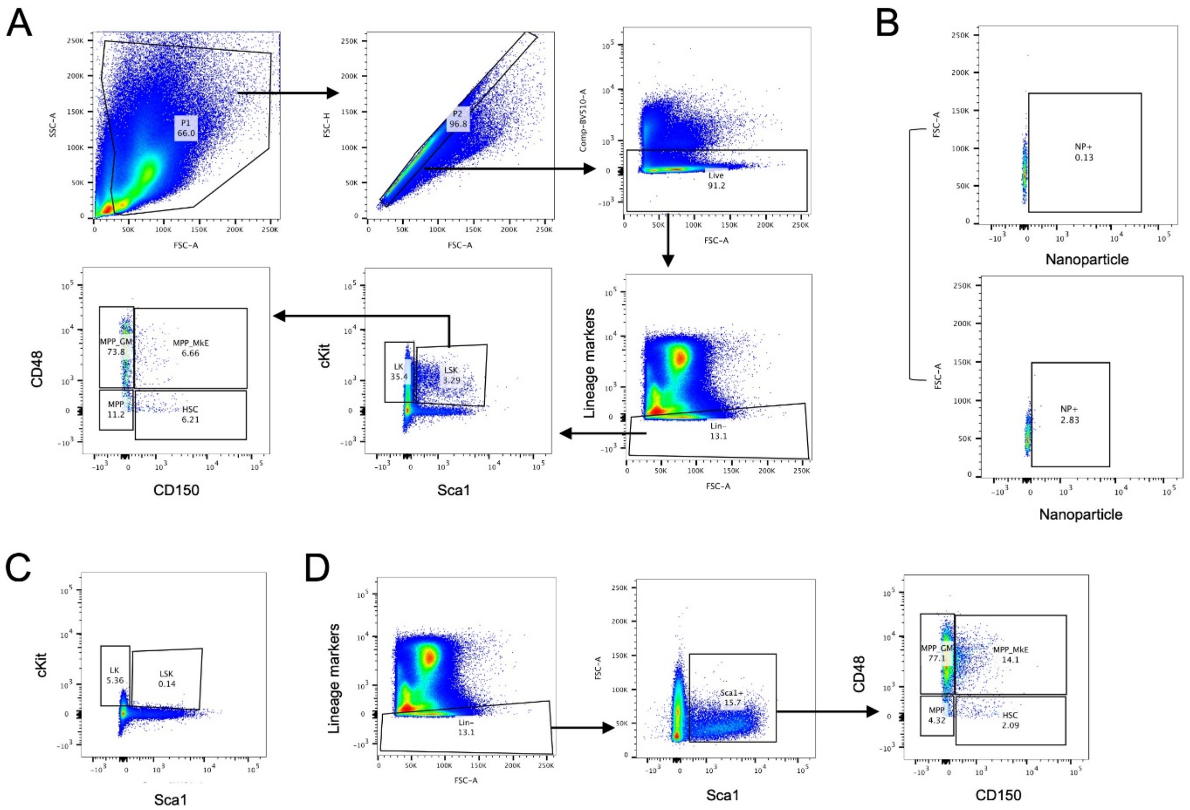


**Supporting Figure 7. Flow gating strategy used for the analysis of bone marrow cells.** (A) Gating strategy based on different surface marker expression. (B) Representative gating for a control sample (no NP signal, top) and a sample with NP signal (bottom). (C) LSK gating obtained in animals that received the cKit–PAA NPs group, where the cKit expression is decreased. (D) Alternative gating strategy for cKit-treated animals, that includes all the Lin- Sca1+ cells.

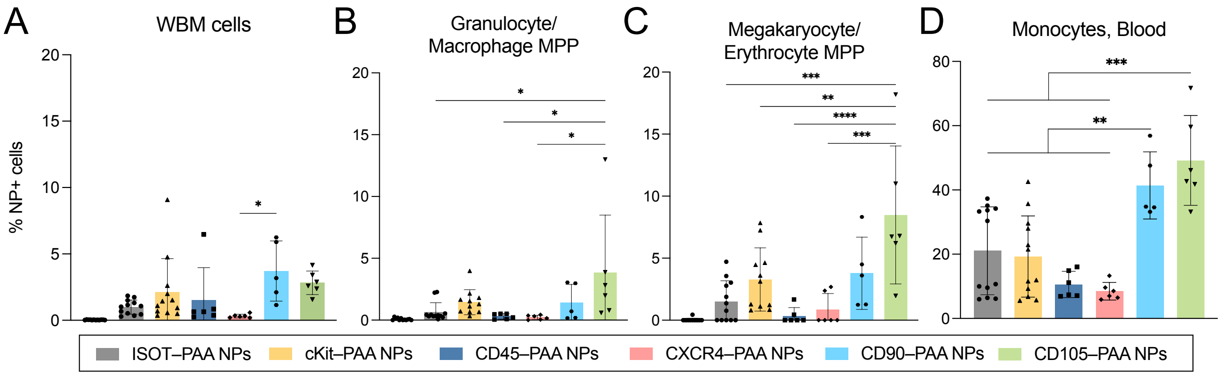


**Supporting Figure 8. Ab–PAA NP targeting capacity to hematopoietic progenitors.** Ab–PAA NP association to (A) whole bone marrow (WBM) cells, (B) Granulocyte/Macrophage MPPs, (C) Megakaryocyte/Erythrocyte MPPs, and (D) blood-circulating monocytes. Statistical analysis is a one-way ANOVA with Tukey correction (*$p<0.05$, **$p<0.01$, ***$p<0.005$, ****$p<0.001$). Bars represent mean, and each dot is one animal replicate ($n \geq 5$ per group).

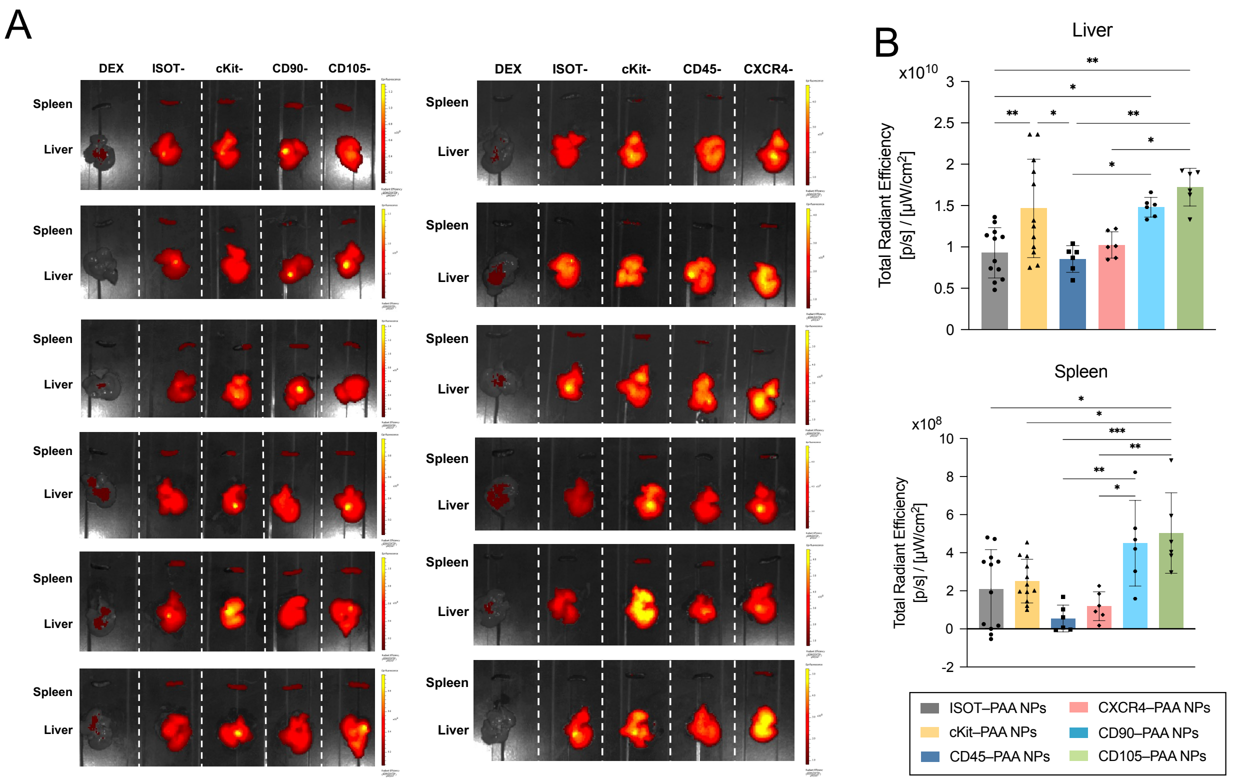


**Supporting Figure 9. IVIS analysis of the organs harvested 1.5 h after intravenous injection of the Ab–PAA NPs.** (A) IVIS images of the fluorescence of the Ab–PAA NPs (Cy5-lipid) signal in the organs collected (liver and spleen) for the different group treatments. (B) Radiant efficiency measurements obtained via IVIS of the fluorophore (Cy5)-tagged Ab–PAA NPs in spleen; bars represent the mean, and each dot represents one biological replicate (n≥6 animals per group). Statistical analysis is a one-way ANOVA with Tukey correction (*p,0.05, **p<0.01, ****p<0.001).

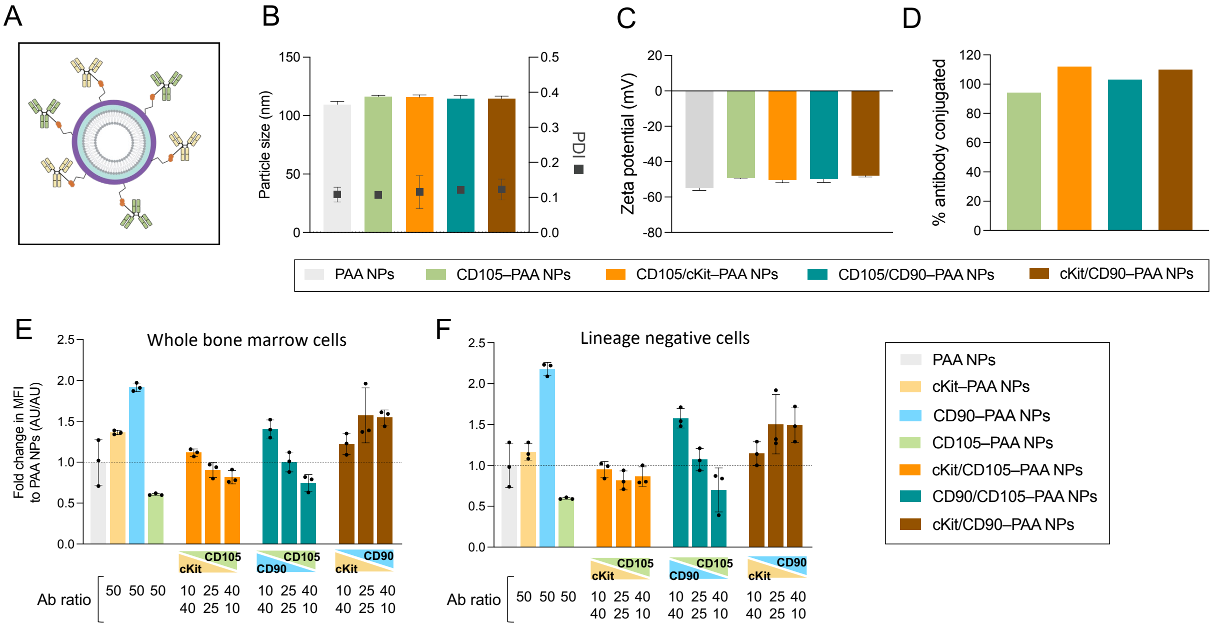


**Supporting Figure 10. Development of dual-targeted PAA NPs.** (A) Schematic of a dual targeted Ab–PAA NP conjugated to two types of Abs through their hinge region to the handles presented on the surface of the outer layer. (B, C) Average particle size (Z-average), polydispersity index (PDI) and ζ-potential of the dual targeted Ab–PAA NPs, in comparison to PAA NPs and CD105–PAA NPs. (D) Yield of the conjugation reaction after purification of the conjugation reaction. Bars represent mean ± standard deviation of 1 technical replicate with 3 measurements. (E, F) Median fluorescence intensity (MFI) values after 1.5 h of incubation with (A) WBM cells, or (B) Lin- cells, normalized to PAA NP, for the combinations of Abs. Statistical analysis is a one-way ANOVA with Tukey correction (***$p<0.005$, ****$p<0.001$). Bars represent mean, and each dot is one technical replicate (≥2 biological replicates for B–C, 1 biological replicate for D-F; each with 3 technical replicates).

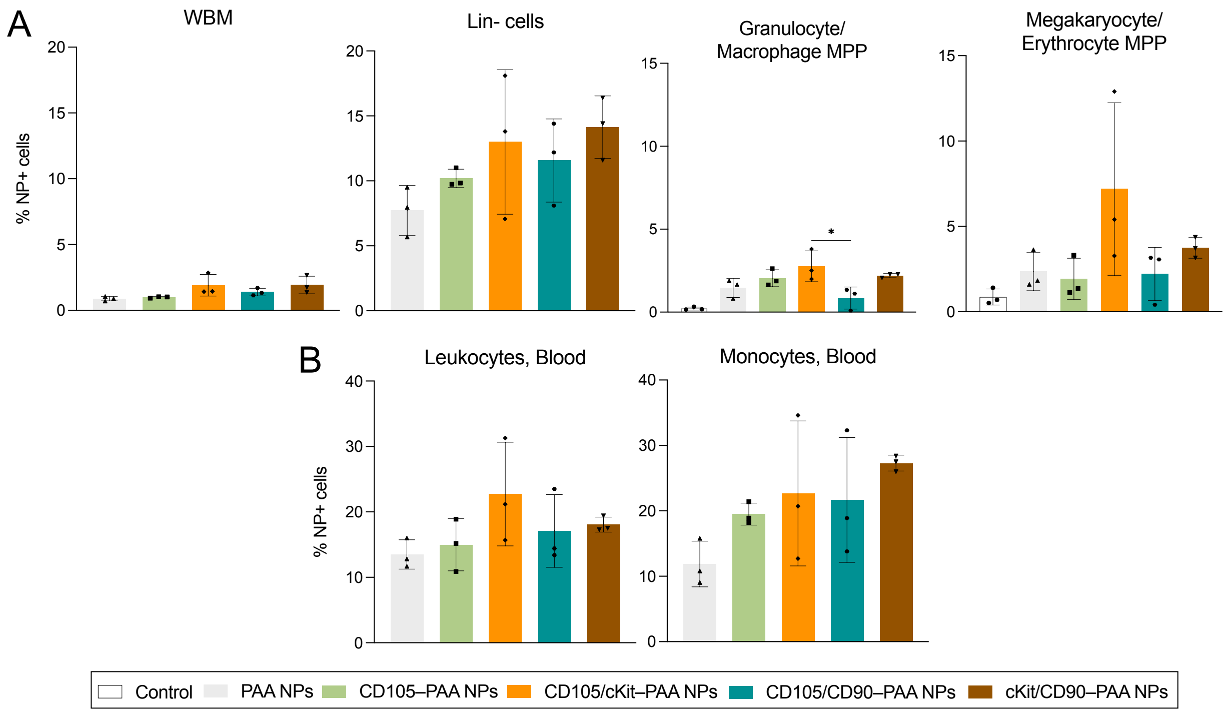


**Supporting Figure 11. HSPC targeting capacity of dual targeted Ab–PAA NPs - after intravenous administration.** (A) Ab–PAA NP association to WBM cells, Lin- cells, Granulocyte/Macrophage MPPs, and Megakaryocyte/Erythrocyte MPPs. (B) Ab–PAA NP-positive cells in blood cells for the different dual targeted Ab–PAA NPs: leukocytes and monocytes. FACS gating strategy identifies the different cell subtypes first, and then the NP-positive populations. Bars represent the mean, and each dot represents one biological replicate (n=3 mice per group). Statistical analysis is a one-way ANOVA with Tukey correction (*p<0.05, **p<0.01, ***p<0.005).

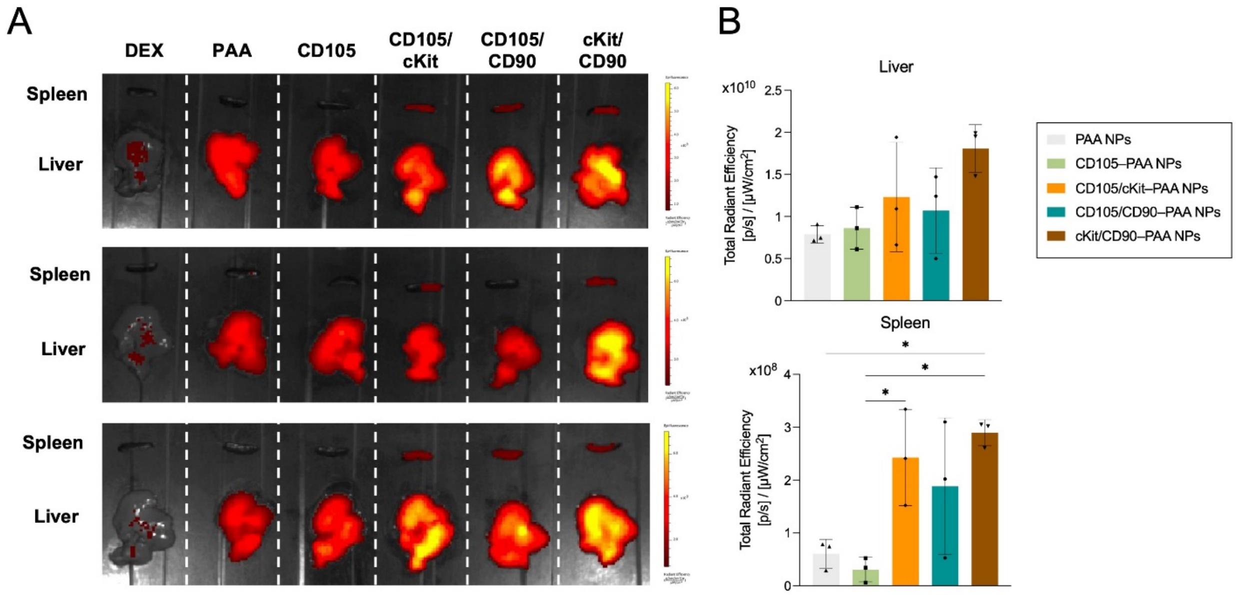


**Supporting Figure 12. IVIS analysis of the organs harvested 1.5 h after intravenous injection of the dual targeted Ab–PAA NPs.** (A) IVIS images of the fluorescence of the nanoparticle (Cy5-lipid) signal in the organs collected (spleen and liver) for the different group treatments. (B) Radiant efficiency measurements obtained via IVIS of the fluorophore (Cy5)-tagged dual targeted Ab–PAA NPs in liver and spleen; bars represent the mean, and each dot represents one biological replicate. Bars represent the mean, and each dot represents one biological replicate (n=3 mice per group). Statistical analysis is a one-way ANOVA with Tukey correction (*p<0.05).

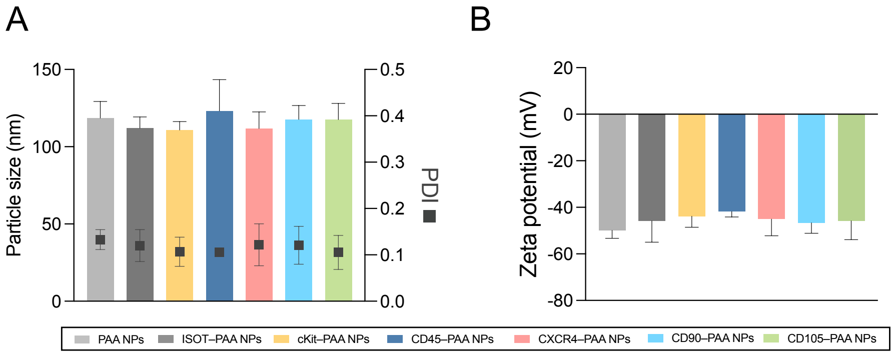


C

| Ab | **Isotype** | **cKit** | **CD45** | **CXCR4** | **CD90** | **CD105** |
|---|---|---|---|---|---|---|
| # linkers | 1.8 ± 0.4 | 1.3 ± 1.2 | 1.9 ± 1.6 | 0.7 ± 0.5 | 0.8 ± 0.5 | 0.7 ± 0.5 |

**Supporting Figure 13. Development of the anti-human Ab–PAA NPs.** (A, B) Average particle size (Z-average), polydispersity index (PDI) and ζ-potential of anti-human Ab–PAA NPs. Bars represent mean ± standard deviation (n≥3 replicates). (C) Table summarizing number of linkers conjugated per Ab (n≥3 replicates).

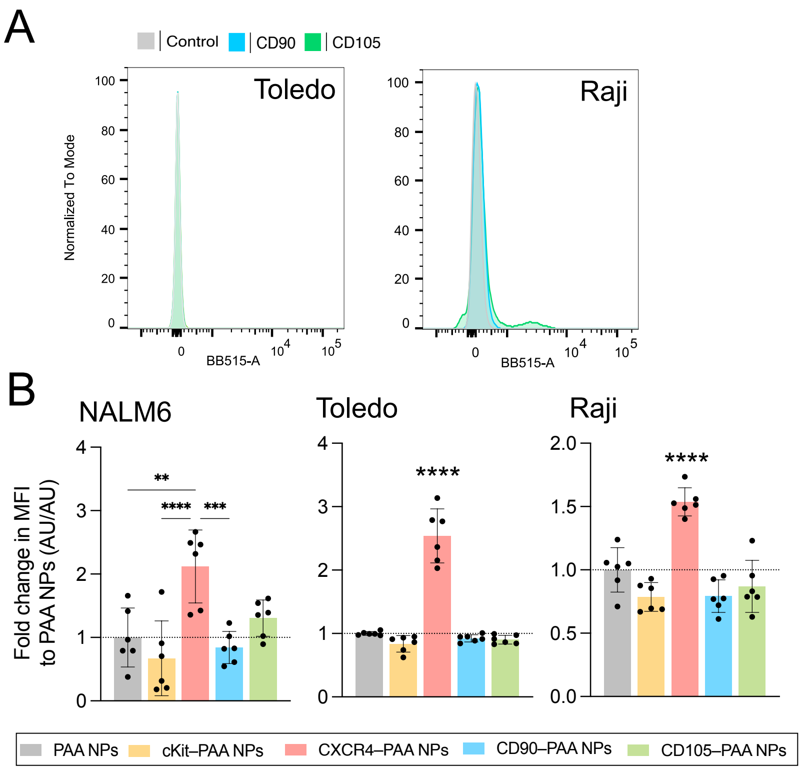


**Supporting Figure 14.** (A) Expression of surface receptors for CD90 and CD105 on Toledo and Raji cells. (B) Median fluorescence intensity (MFI) values after 4 h of incubation for anti-human Ab–PAA NPs, normalized to PAA NPs, in NALM6 cells, Toledo cells, and Raji cells. Statistical analysis is a one-way ANOVA with Tukey correction (**p<0.01, ***p<0.005, ****p<0.001). Bars represent mean with standard deviation; each dot is one technical replicate (2 biological replicates with 3 technical replicates each).
